# Transcriptomic and morphological response of SIM-A9 mouse microglia to carbon nanotube neuro-sensors

**DOI:** 10.1101/2020.06.30.181420

**Authors:** Darwin Yang, Sarah J. Yang, Jackson Travis Del Bonis-O’Donnell, Rebecca L. Pinals, Markita P. Landry

## Abstract

Single-walled carbon nanotubes (SWCNT) are used in neuroscience for deep-brain imaging, neuron activity recording, measuring brain morphology, and imaging neuromodulation. However, the extent to which SWCNT-based probes impact brain tissue is not well understood. Here, we study the impact of (GT)_6_-SWCNT dopamine nanosensors on SIM-A9 mouse microglial cells and show SWCNT-induced morphological and transcriptomic changes in these brain immune cells. Next, we introduce a strategy to passivate (GT)_6_-SWCNT nanosensors with PEGylated phospholipids to improve both biocompatibility and dopamine imaging quality. We apply these passivated dopamine nanosensors to image electrically stimulated striatal dopamine release in acute mouse brain slices, and show that slices labeled with passivated nanosensor exhibit higher fluorescence response to dopamine and measure more putative dopamine release sites. Hence, this facile modification to SWCNT-based dopamine probes provides immediate improvements to both biocompatibility and dopamine imaging functionality with an approach that is readily translatable to other SWCNT-based neurotechnologies.

## Introduction

Nanoscale neurotechnologies often demonstrate increased biocompatibility and less invasive implementation than their micro- or macro-scale counterparts (*1*), and can offer higher signal-to-noise ratios owing to the relatively large surface area of nanoscale materials (*2*). For this reason, engineered nanoparticles have recently demonstrated broad-scale utility in neuroscience for neurological recordings (*3*), drug delivery across the blood-brain barrier (*4*), and for brain imaging (*5*). In particular, carbon nanotubes have shown increasing applicability for neuron stimulation, electrochemical recordings of neuron action potentials, mapping brain extracellular space, deep-brain imaging, and imaging neurotransmission (*6–12*). Towards the last point, recent developments are enabling imaging of chemical communication between cells, specifically for a class of neurotransmitters known as neuromodulators whose imaging has eluded existing methods of inquiry (*13*). Previous work includes development of a nanoscale near infrared catecholamine probe, nIRCat, that can capture dopamine release and reuptake kinetics in the brain striatum, and measure the influence of drugs on these signaling properties (*11*). nIRCat is synthesized by noncovalent conjugation of (GT)_6_ single stranded DNA (ssDNA) and near infrared (nIR) fluorescent single-walled carbon nanotubes (SWCNTs) (*14*). However, as these and numerous other nanoscale neurotechnologies based on SWCNTs are used to probe the brain microenvironment, it becomes important to understand how these nanoparticles affect surrounding brain tissue.

Carbon nanomaterials have previously been implicated in activation of the innate immune system across multiple biological organisms and through numerous mechanisms. Nonspecific adsorption of complement proteins in serum causes recognition of carbon nanotubes and activation of the complement system (*15*). In macrophages, interaction of carboxylic acid functionalized SWCNTs with toll-like receptors 2 and 4 (TLR2/4) results in activation of an inflammatory signaling cascade and protein expression of cytokines (*16*). Release of cytokines such as interleukin 1β (*Il1b*) then propagates the inflammatory response to surrounding tissue. However, the biological impact of carbon nanomaterials—particularly those with pristine graphene lattices—has not been well characterized in the brain. Inflammation in the brain has long been associated with multiple negative health outcomes including neurotoxicity, neurodegeneration, and loss of function (*17, 18*). These effects are particularly consequential in the context of studying chemical neurotransmission. Therefore, it is imperative to characterize and quantify the extent to which carbon nanotubes induce an inflammatory response in the brain and if such effects can be mitigated.

Microglia are specialized immune cells found in the central nervous system. Recognition of tissue damage or pathogenic material causes microglial activation, characterized by a change in cell morphology and an inflammatory response. This response promotes clearance of the pathogen through phagocytosis, and has been shown to result in neurotoxicity and reduced dopamine concentrations in the striatum (*19, 20*). Larger multi-walled carbon nanotubes with carboxylic acid functionalization have previously been found to negatively impact microglial phagocytosis processes (*21, 22*). Therefore, probing the impact of carbon nanotubes on microglia is of critical importance to assess the biocompatibility of SWCNT-based neuro-technologies.

In this work, we study the transcriptomic effects induced by SWCNT dopamine nanosensors on SIM-A9 microglia. The SIM-A9 cell line was spontaneously immortalized from primary mouse microglia, and exhibits similar characteristics as primary microglia including morphology, response to endotoxin exposure, and cytokine secretion (*23*), making this cell line optimal to study neuroinflammation (*24, 25*). We quantify microglial morphological and transcriptomic responses induced by SWCNTs compared to those elicited by exposure to other probes commonly used in neuroscience, including calcium and voltage-sensitive probes, and adeno-associated viral (AAV) vectors (*26*). Specifically, we utilize live-cell imaging, quantitative PCR, and RNA-seq to elucidate and quantify the morphological and inflammatory cell mechanisms affected by exposure to these probes. We then use knowledge of these responses to develop a method to passivate SWCNT dopamine nanosensors with a polyethylene glycol (PEG) conjugated phospholipid to improve nanosensor biocompatibility and mitigate attenuation of nanosensor efficacy when used for in-brain imaging. Finally, we show that this passivation methodology improves nanosensor dopamine imaging in excised mouse brain tissue with electrically evoked neurotransmitter release.

## Results & Discussion

### Morphological response of SIM-A9 microglia to SWCNTs

We first studied the effects of the (GT)_6_-SWCNT catecholamine nanosensor on SIM-A9 microglial cell morphology, a phenotypic marker of microglial activation that promotes cell migration. Specifically, a morphology change from round to ramified is characteristic of microglial activation *in vitro* (*27*). Incubation of SIM-A9 microglia with 5 μg/mL (GT)_6_-SWCNTs resulted in drastic cell morphology change within 4 h post-exposure. Cells progressed from round, amoeboid morphologies to highly branched, ramified structures displaying elongated cellular processes (Fig. 1A), while control cells absent from exposure to (GT)_6_-SWCNTs retained a round morphology (Fig. 1B). Live-cell imaging time lapse videos show control SIM-A9 populations consisted of round, highly motile cells (movie S1). Incubation of cells with (GT)-SWCNTs caused immediate ramification of SIM-A9 microglia and loss of cell motility during the first two hours post exposure (movie S2).

**Figure 1.**
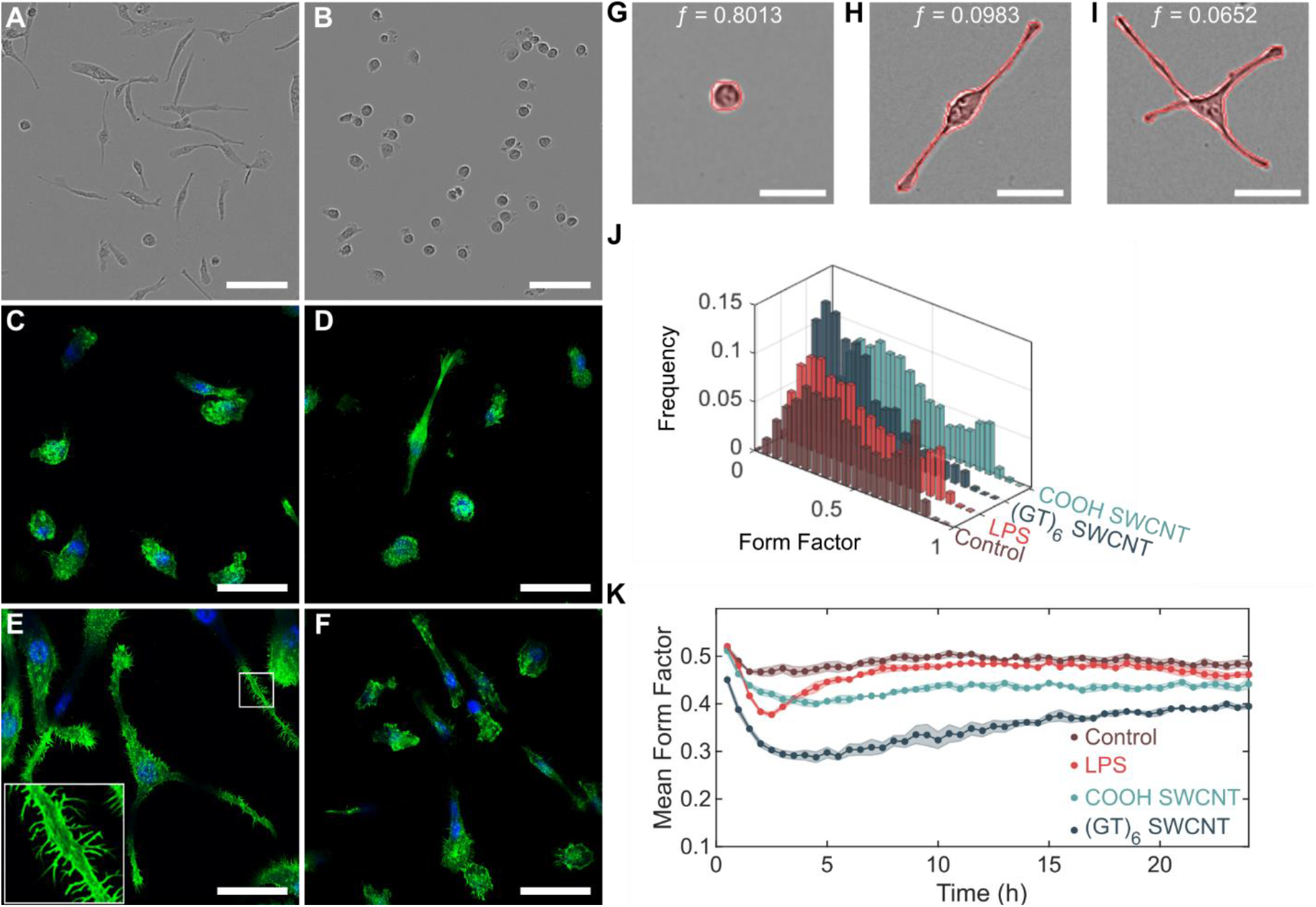
SWCNT-induced morphology change in SIM-A9 microglial cells. **(A and B)** Phase contrast images of SIM-A9 microglia following 4 h incubation with **(A)** 5 μg/mL (GT)_6_-SWCNT or **(B)** PBS. Scale bars are 100 μm. **(C to F)** Confocal fluorescence microscopy images of FAM-stained F-actin (green) of fixed microglia following 4 h incubation with **(C)** PBS, **(D)** 5 ng/mL LPS, **(E)** 5 μg/mL (GT)_6_-SWCNT, and **(F)** 5 μg/mL COOH-SWCNT. Nuclei are counterstained with DAPI (blue). Scale bars are 50 μm. **(G to I)** SIM-A9 morphologies and corresponding form factor values for **(G)** round, **(H)** bipolar, and **(I)** multipolar cells. Outlines of the identified cells are shown in red. Scale bar is 50 μm. **(J)** Form factor distribution following 3 h incubation with PBS (control), 5 ng/mL LPS, 5 μg/mL (GT)_6_-SWCNT, and 5 μg/mL COOH-SWCNT. **(K)** Tracking of mean form factor per field of view capture over 24 h. Shaded regions represent standard error of the mean (N=3).

Cell morphology change in the (GT)_6_-SWCNT-treated cells coincided with actin cytoskeletal growth as measured with F-actin probe phalloidin conjugated with Alexa Fluor 488, forming projections resembling microglial filopodia (Fig. 1, C to F). These projections are known to be responsible for increasing microglial cell surface area within the brain microenvironment as a result of microglial activation and are typically found at the tips of microglial processes (*28, 29*). Conversely, we observe that (GT)_6_-SWCNT exposure promoted growth of projections along the entire length of the cell branches, not only at the tip of microglial processes (Fig. 1E). Interestingly, positive control experiments of SIM-A9 cells incubated with lipopolysaccharide (LPS), a class of molecule found in gram-negative bacterial cell walls known to activate toll-like receptor 4 (TLR4) and induce a strong inflammatory response, induced a relatively marginal change in cell morphology compared to (GT)_6_-SWCNT exposure. Hence, the pathway of microglial activation by SWCNTs may be distinct from previously observed TLR4 activation by carbon nanomaterials (*16*). Carboxylic acid functionalized SWCNTs (COOH-SWCNTs) were included as an additional positive control, and similarly induced a marginal change in cell morphology compared to (GT)_6_-SWCNTs. These latter results suggest a strong influence of nanomaterial surface chemistry on nanoparticle biocompatibility.

Cell morphology change in time lapse videos was quantified by assigning each cell with a form factor value computed by the following equation:

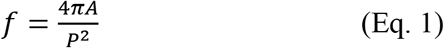

where A is the area occupied by a cell and P is the perimeter of the cell. Form factor values near 1 therefore indicate round cells (Fig. 1G), whereas decreasing values of *f* correlate with higher degrees of ramification (Fig. 1, H to I). Untreated SIM-A9 cells reveal a defined population of cells with form factors near 0.8 (Fig. 1J). Following a 3 h incubation of SIM-A9 cells with 5 ng/mL LPS or 5 μg/mL COOH-SWCNTs, this *f* = 0.8 peak is diminished, whereas in cells treated with 5 μg/mL (GT)_6_-SWCNTs, the peak at *f* = 0.8 is not observed and instead skews to lower form factor values. To determine the time-dependence of this morphology change, we performed live-cell imaging of the above samples and averaged the form factor values of all cells within a given field of view to track progression of mean cell form factor for 24 h post-treatment. Following addition of (GT)_6_-SWCNTs, an immediate decrease in mean form factor occurred within 1 h (Fig. 1K). Presence of (GT)_6_-SWCNTs caused mean form factor to decrease to a minimum of 0.288 ± 0.011 at 4.5 h post-exposure, compared to a minimum value of 0.473 ± 0.010 for untreated control cells. LPS-stimulated cells reached a minimum mean form factor of 0.377 ± 0.004 at 2.5 h post-exposure and returned to near baseline levels after approximately 7.5 h. Conversely, SWCNT-stimulated cells failed to return to morphologies consistent with the control cell population within 24 h. The mean form factor of (GT)_6_-SWCNT treated SIM-A9 24 h post-exposure was 0.395 ± 0.005 compared to 0.483 ± 0.10 for untreated control cells. The extent of cell ramification caused by (GT)_6_-SWCNTs was concentration-dependent from 0.1 to 5 μg/mL (fig. S1A), coinciding with a SWCNT concentration range relevant for neuro-applications (*11*). To confirm that the observed morphology changes were due to SWCNTs and not the (GT)_6_ oligonucleotide alone, we imaged SIM-A9 cells exposed to (GT)_6_-_ss_DNA. We found that 1.67 μM (GT)_6_-_ss_DNA oligonucleotides alone did not cause a significant change in SIM-A9 cell morphology at any time point within a 24 h live-cell imaging experiment (fig. S1B). This concentration corresponds to the total DNA concentration in a 10 μg/mL (GT)_6_-SWCNT suspension, further suggesting that the above-discussed effects are induced by the SWCNT carbon lattice.

5 μg/mL (GT)_6_-SWCNTs were observed to internalize into SIM-A9 cells within 1 h of incubation at 37°C, 5% CO_2_ (fig. S2A). We find that internalization is energy-dependent, as observed by the absence of SWCNT internalization in SIM-A9 cells at 4°C (fig. S2B). Previous studies of SWCNT internalization in mammalian cells have determined the internalization mechanism to be predominantly energy-dependent clathrin-mediated endocytosis (*30, 31*). No correlation was found between degree of internalization and cell morphology at 2 h post exposure to SWCNTs (fig. S2, C and D), suggesting that the cellular morphological change is due to cell signaling rather than a physical interaction between SWCNTs and actin filaments. Furthermore, we find that phagocytosis of fluorescent Zymosan particles by SIM-A9 cells was diminished following exposure to concentrations greater than or equal to 0.5 μg/mL SWCNTs (fig. S3), where reduced phagocytosis is characteristic of quiescent, ramified microglia in rats, coinciding with a morphology change (*32*).

### SIM-A9 microglia transcriptomic response to neuro-probes

We utilized high throughput mRNA sequencing to determine and quantify the full transcriptomic response of SIM-A9 microglia cells to 10 μg/mL (GT)_6_-SWCNT exposure. This response was compared to that induced by 10 μg/mL COOH-SWCNT, 10 ng/mL LPS positive control, and other commonly used neuro-probes including those for calcium imaging (2 μM Fura-2), voltage sensing (2 μM DiSBAC_2_ and Di-2-ANEPEQ), and AAV viral vector (50,000 virus molecules per SIM-A9 cell). Concentrations of these molecular probes were chosen to be reflective of their working concentrations for brain imaging applications. The non-SWCNT neuro-probes screened did not induce a noticeable morphology change in SIM-A9 microglia (fig. S1C). Furthermore, multidimensional scaling (MDS) analysis of the normalized gene counts for each sequencing library revealed close clustering of biological replicates of Fura-2, DiSBAC_2_, and Di-2-ANEPEQ with the untreated microglia control (Fig. 2A), suggesting these small molecule neuro-probes have a minor impact on microglial cell function over 2 h. AAV sequencing libraries also did not show significant divergence from control at the 2 h time point. However, MDS analysis showed deviation of SWCNT and LPS incubated cell samples from the untreated SIM-A9 control. Hierarchical clustering of sequencing libraries further demonstrated that cells incubated with Fura-2, DiSBAC(2), Di-ANEPEQ, and AAV did not elicit a significant transcriptomic response, evidenced by the statistical similarity of these sequencing libraries to untreated control libraries (Fig. 2B). Hence, downstream differential gene expression analysis and ontological analysis was only carried out at a dendrogram cut height of 30, comparing SIM-A9 cells treated with (GT)_6_-SWCNTs, COOH-SWCNTs, and LPS to untreated control groups.

**Figure 2.**
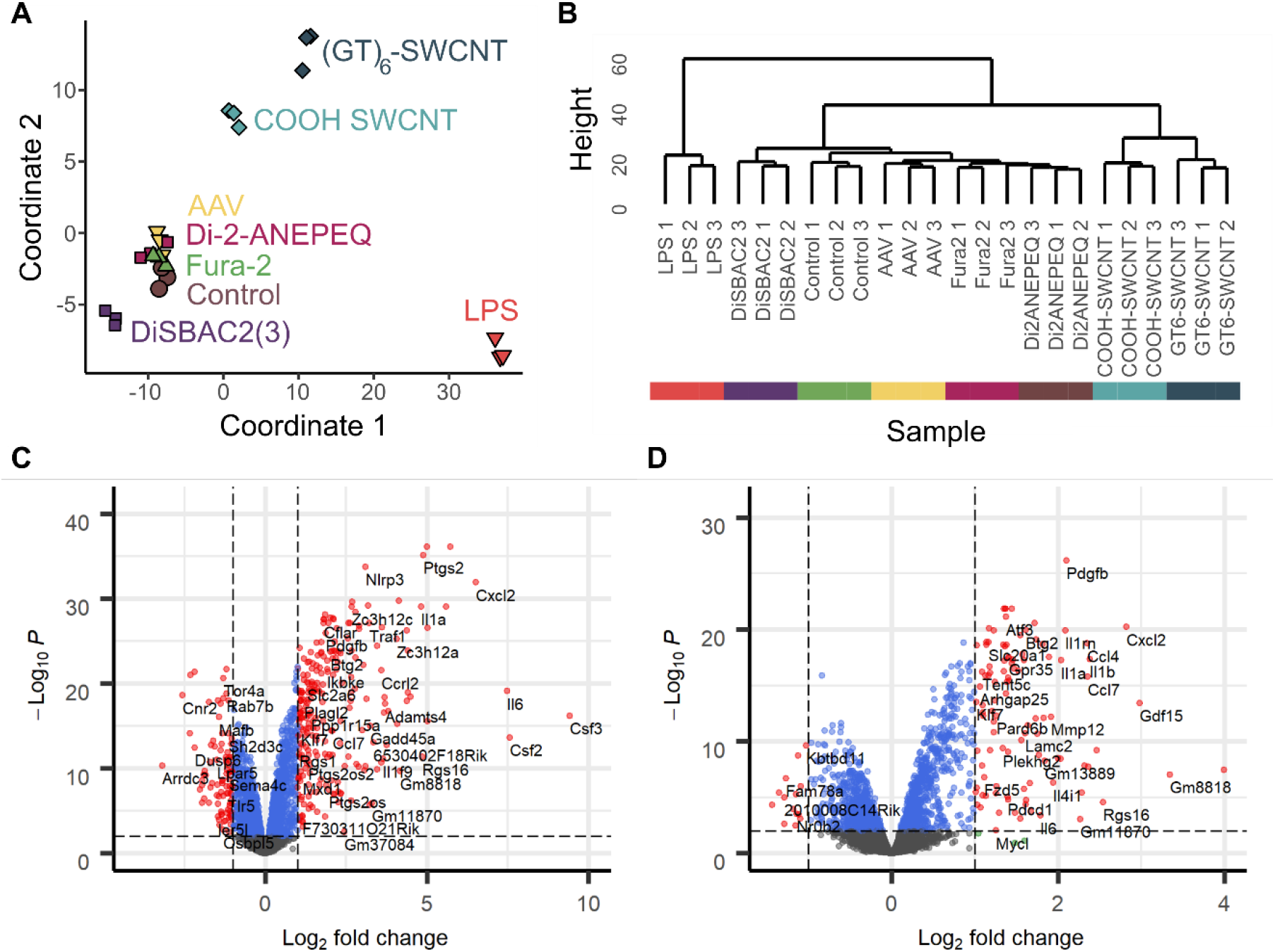
Transcriptomic response of SIM-A9 microglia to neuro-probes. **(A)** Multidimensional scaling analysis of gene count tables generated for each RNA-seq library, where each marker represents a biological replicate. Axes represent the two principal components of highest variance. Control represents untreated SIM-A9 microglial cells. **(B)** Hierarchical clustering of sequenced libraries based on normalized gene counts. **(C and D)** Volcano plots of **(C)** LPS and **(D)** (GT)_6_-SWCNT incubated SIM-A9 cells showing log_2_ fold change in gene expression vs. log_10_ adjusted p value for all 9770 identified genes, relative to untreated control cells. Horizontal and vertical dashed lines delineate p_adj_ = 0.05 and log_2_ fold change = 1 respectively.

We performed differential gene expression analysis using the *edgeR* package (*33*). SIM-A9 cell experimental groups were compared pairwise to untreated control groups. For LPS-treated samples, out of 9770 genes identified across all sequencing libraries, 332 were both differentially expressed (p_adj_ < 0.05) and exhibited a greater than 2-fold change in expression vs. the untreated control (table S1, Fig. 2C), where p_adj_ is the false discovery rate corrected p value. Only 119 such genes were identified for (GT)_6_-SWCNT vs. the untreated control, and 49 genes for COOH-SWCNT vs. untreated control. LPS promoted upregulation of many inflammatory cytokines such as *Csf2, Csf3, Il1b*, and *Cxcl2* (Fig. 2C), where the latter two cytokines are also among the most highly upregulated genes by (GT)_6_-SWCNTs (Fig. 2D). However, the SWCNT-induced expression-fold change for these genes was significantly lower than that caused by LPS.

Conversely, platelet-derived growth factor subunit B, *Pdgfb*, was more significantly upregulated by (GT)_6_-SWCNTs than by LPS, and also shows highly statistically significant upregulation by COOH-SWCNTs (fig. S4A), thus is a potential biomarker for cellular response to SWCNTs.

The *topGO* R package was used to perform enrichment analysis on differentially expressed genes with a cutoff of p_adj_ < 0.01 (*34, 35*). Specifically, we examined enrichment of Gene Ontology (GO) biological processes terms. As expected, differentially expressed genes in LPS vs. untreated control groups showed high overrepresentation of processes associated with toll-like receptor signaling and inflammation, including *Cellular response to lipopolysaccharide* (Fig. 3A). Exposure of SIM-A9 cells to (GT)_6_-SWCNTs caused enrichment of similar inflammatory GO terms, including the most statistically significantly enriched GO term, *Inflammatory response* (Fig. 3B). Both LPS and (GT)_6_-SWCNT treatments caused enrichment of *Positive regulation of ERK1 and ERK2 cascade* GO term. Of 97 annotated genes, 52 were differentially expressed in LPS samples out of an expected 24.4, giving a gene set enrichment p value of 3.7 x 10^-9^. (GT)_6_-SWCNT treatment caused differential expression of 41 annotated genes of an expected 16.2, with a corresponding p value of 5.4 x 10^-9^. Enrichment of ERK signaling by both LPS and (GT)_6_-SWCNTs may indicate similar inflammatory signal transduction induced by the two molecules.

**Figure 3.**
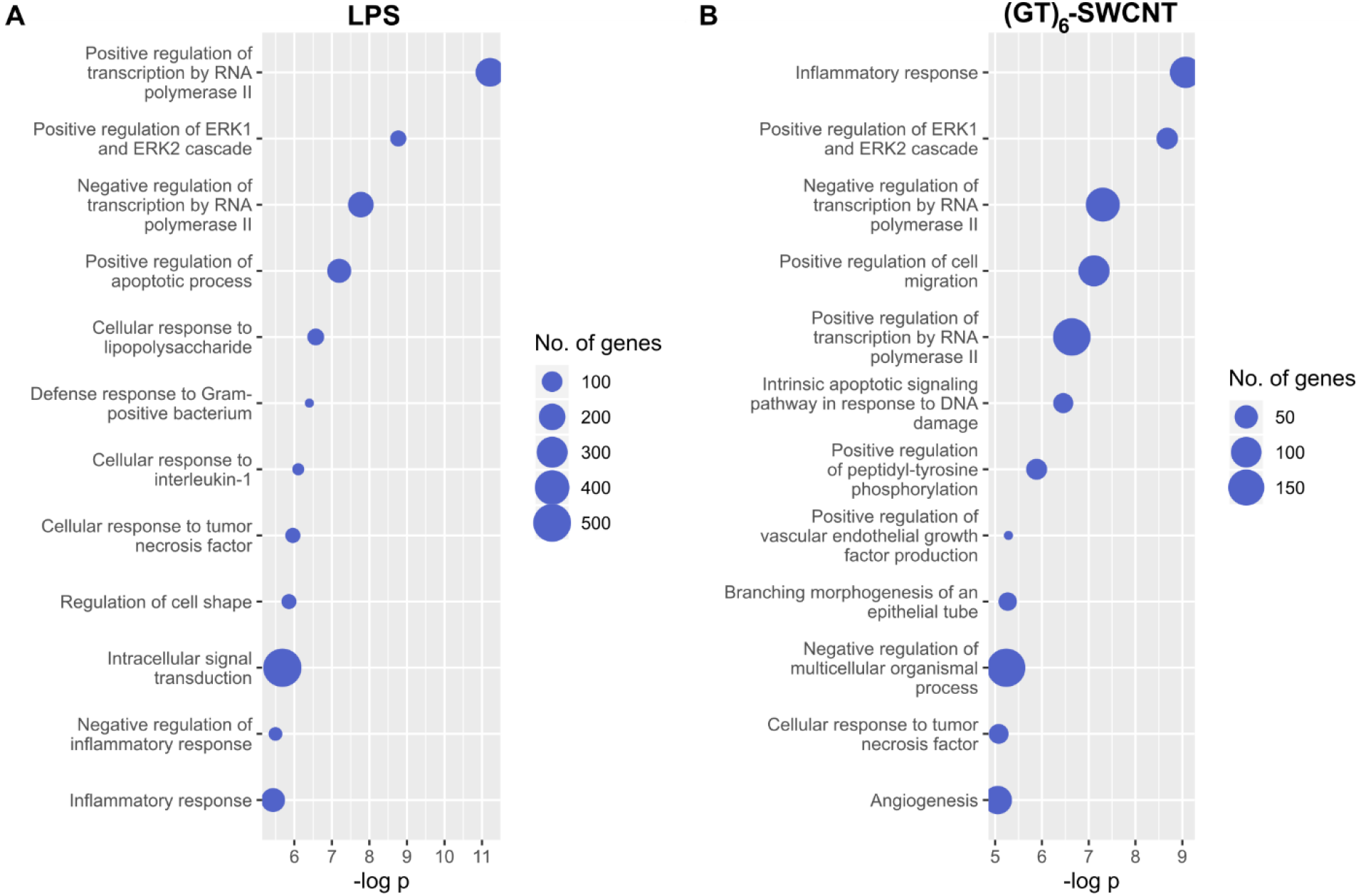
Gene Ontology enrichment analysis of LPS and (GT)_6_-SWCNT treated SIM-A9 microglia. **(A and B)** Overrepresentation analysis of differentially expressed genes identified in RNA sequencing libraries for SIM-A9 microglia stimulated with **(A)** LPS and **(B)** (GT)_6_-SWCNT. Top twelve most highly enriched ontologies are displayed.

Other biological processes overrepresented by exposure of microglia to (GT)_6_-SWCNT samples include GO terms related to tissue development such as *Branching morphogenesis of an epithelial tube* and *Angiogenesis*, where the nonspecific nature of these processes in relation to microglial cells may suggest noncanonical activation of cellular mechanisms. Lastly, similar to (GT)_6_-SWCNTs, COOH-SWCNTs promoted enrichment of ERK signaling terms (fig. S4B), pointing to the significance of this signaling cascade in SWCNT-induced immune responses. Interestingly, COOH-SWCNTs caused differential expression of a larger number of pseudogenes in SIM-A9 microglia than either LPS or (GT)_6_-SWCNTs (table S1), despite eliciting only a minor morphological change (Fig.1, J and K). This may be due to overrepresentation of GO terms in the set of COOH-SWCNT induced differentially expressed genes, including *Regulation of transcription* (fig. S4B).

### Passivation of ssDNA-SWCNT neuro-sensors with phospholipid-PEG

A common approach to improving biocompatibility of nanotechnologies involves nanoparticle surface functionalization with polyethylene glycol (PEG) polymers to promote steric exclusion of proteins, increase nanoparticle hydrophilicity, and thereby prevent subsequent immune activation. However, covalent linkage of molecules to the SWCNT carbon lattice has been shown to abate SWCNT fluorescence required for neuro-imaging applications (*36*). Consequently, we developed a noncovalent modification strategy for passivation of (GT)_6_-SWCNT nanosensors with PEGylated phospholipids that display a high affinity for the SWCNT surface. SWCNTs have previously been dispersed using PEGylated phospholipids to form highly disperse suspensions (*37, 38*), however, the creation of a hybrid ssDNA and PEG-phospholipid SWCNT surface coatings for dual sensing and biocompatibility purposes remains unexplored. We used saturated 16:0 PEG-phosphatidylethanolamines (PEG-PE) with varying PEG molecular weights ranging from 750 Da to 5000 Da to form co-suspensions with (GT)_6_-SWCNTs, then assessed their effect on nanosensor biocompatibility and efficacy. Sonication of (GT)_6_-SWCNTs with PEG_2000_-PE at a 1:1 SWCNT to phospholipid mass ratio caused a decrease in SWCNT nIR fluorescence intensity and a red shifting of the fluorescence emission (Fig. 4B), indicating an increase in the polarity of the SWCNT dielectric environment, consistent with biomolecular adsorption phenomena (*39–41*). This result is recapitulated in the absorbance spectra of (GT)_6_-SWCNTs, where nIR absorbance peaks corresponding to SWCNT E11 transitions are red-shifted upon passivation with PEG-PE with variable PEG molecular weights (fig. S5). The 750 Da PEG phospholipid caused the highest magnitude wavelength shift, whereas larger 2000 Da and 5000 Da PEG phospholipids induced intermediate red-shifting. This larger solvatochromic shift may indicate higher surface density of PEG_750_-PE on the SWCNT surface compared to larger PEG chains.

**Figure 4.**
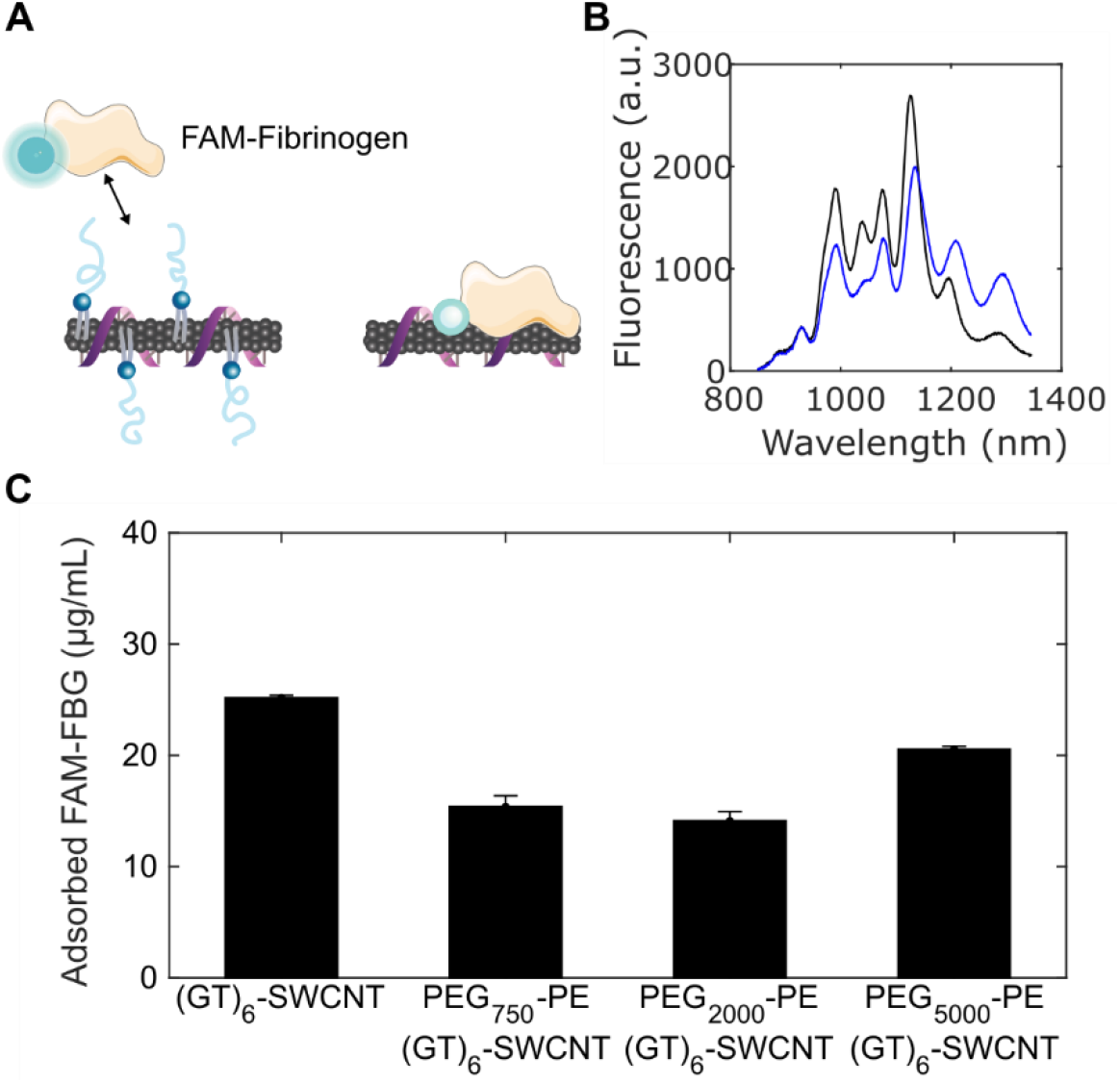
Passivation of (GT)_6_-SWCNTs with PEG-PE phospholipid. **(A)** Schematic of PEG-PE adsorption to (GT)_6_-SWCNTs and subsequently deterring FAM-fibrinogen adsorption. **(B)** Comparison of nIR fluorescence spectra of 5 μg/mL (GT)_6_-SWCNT (black) vs. PEG_2000_-PE/(GT)_6_-SWCNT (blue). **(C)** Concentration of adsorbed FAM-FBG on 5 μg/mL (GT)_6_-SWCNT. Initial concentration of FAM-FBG added to solution was 40 μg/mL. Error bars represent standard error of the mean (N=3).

Nonspecific protein adsorption was quantified on PEG-PE/(GT)_6_-SWCNT passivated nanosensor constructs using a previously developed method for tracking biomolecular adsorption on nanoparticle surfaces in real-time (*41*). Blood coagulation protein fibrinogen (FBG) was selected as a representative binding protein owing to its high affinity for the SWCNT surface (*41*). SWCNT-induced quenching of the fluorophore fluorescein conjugated to fibrinogen (FAM-FBG) was used to determine the degree of adsorption of 40 μg/mL FAM-FBG to 5 μg/mL (GT)_6_-SWCNTs with and without PEG-PE passivation (Fig. 4C, fig. S6, A and B). All molecular weight PEG-PEs caused a reduction in total concentration of adsorbed FAM-FBG after 1 h incubation. Phospholipids with a PEG molecular weight of 2000 Da best mitigated against protein adsorption, showing a 28 ± 2% reduction in adsorption of FAM-FBG after 1 h compared to unmodified (GT)_6_-SWCNT nanosensors. The degree of FAM-FBG adsorption on PEG_2000_-PE/(GT)_6_-SWCNTs was comparable to SWCNTs suspended with solely PEG_2000_-PE (fig. S6C).

We next tested the interaction of PEG_2000_-PE/(GT)_6_-SWCNTs nanosensors with SIM-A9 microglia. Analogous to our protein adsorption mitigation results, the 2000 Da PEG length showed the greatest mitigation in SIM-A9 morphology change (Fig. 5, A to C). Unmodified (GT)_6_-SWCNTs caused mean form factor to decrease to a minimum of 0.490 ± 0.013, whereas the PEG_2000_-PE/(GT)_6_-SWCNTs merely led to a minimum of 0.618 ± 0.005. PEG-PE modified samples also exhibited a return to baseline morphology returning to untreated control levels after 9, 6 and 15 h respectively for 750, 2000, and 5000 Da PEG molecular weights. Conversely, unmodified (GT)_6_-SWCNT nanosensors did not show a return to baseline morphology within the 24 h experiment.

**Figure 5.**
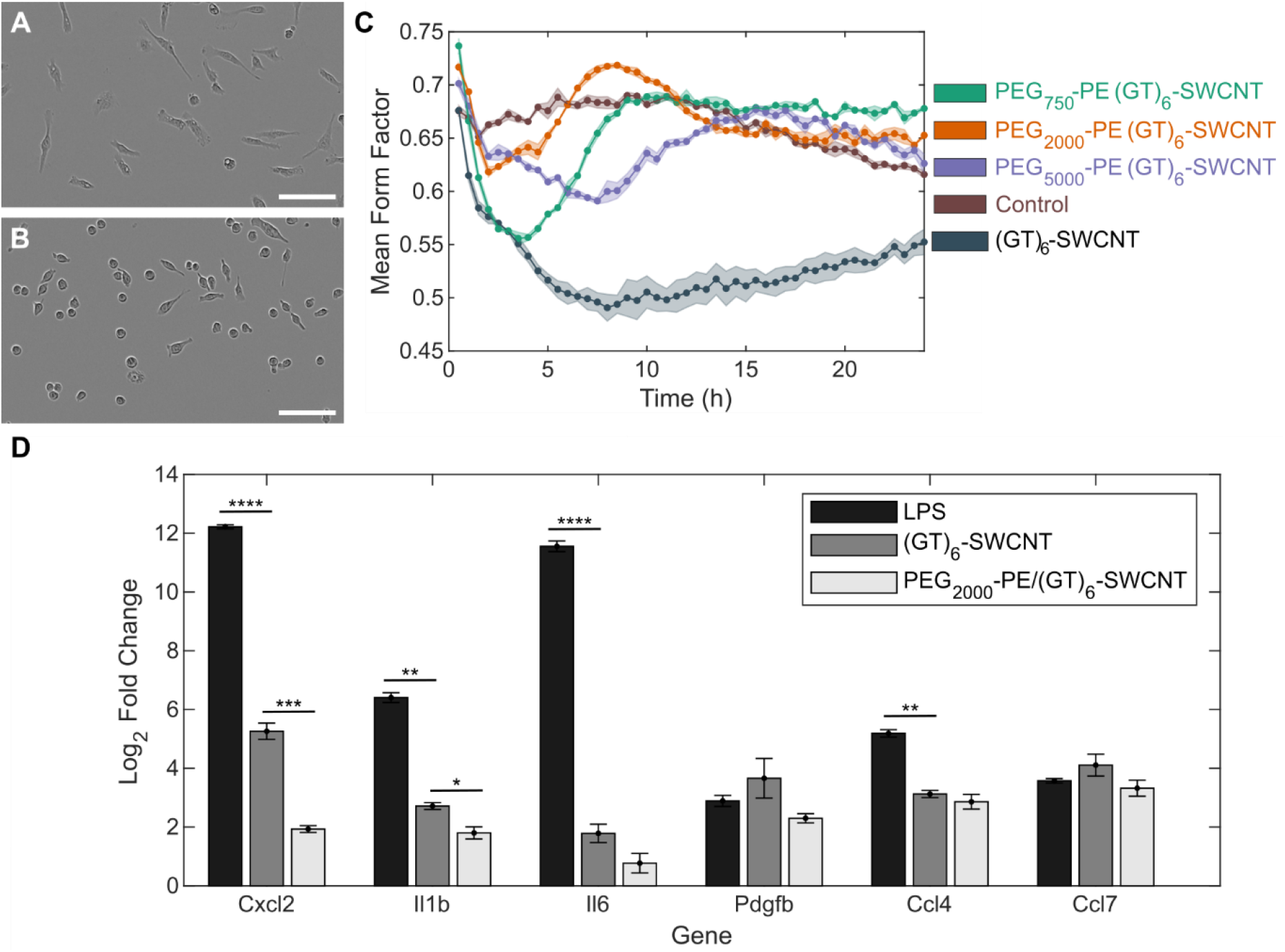
Effect of PEG-PE phospholipid passivation on microglial activation. **(A and B)** Phase contrast images of SIM-A9 microglial cells incubated with **(A)** (GT)_6_-SWCNTs and **(B)** PEG_2000_-PE passivated (GT)_6_-SWCNTs at 5 μg/mL for 6 h. **(C)** Mean form factor time traces of SIM-A9 microglia incubated with PEG-PE passivated vs. unpassivated (GT)_6_-SWCNTs compared to no treatment control. Shaded regions represent standard error of the mean (N=3). **(D)** Quantitative reverse transcriptase PCR expression-fold change of SWCNT inflammatory cytokine markers upon stimulation with 5 ng/mL lipopolysaccharide, 5 μg/mL (GT)_6_-SWCNT, and 5 μg/mL PEG_2000_-PE/(GT)_6_-SWCNT. Incubation time was 2 h. Error bars represent standard error of the mean (N=3). Statistical analyses compare LPS and PEG_2000_-PE/(GT)_6_-SWCNT relative expression changes to that of (GT)_6_-SWCNT: * p < 0.05, ** p < 0.005, *** p < 5 x 10^-4^, **** p < 5 x 10^-5^.

We next tested the inflammatory response induced by PEG_2000_-PE/(GT)_6_-SWCNT nanosensors exposed to SIM-A9 microglia. Noncovalent passivation of ssDNA-wrapped SWCNTs with PEG-PE phospholipid caused a reduction in SIM-A9 inflammatory response, exemplified by a decrease in the expression of inflammatory cytokines previously identified as upregulated in our transcriptomic studies. Specifically, genes *Cxcl2, Il1b, Il6, Pdgfb, Ccl4*, and *Ccl7* were selected as SWCNT-specific upregulated biomarkers from the (GT)_6_-SWCNT and COOH-SWCNT libraries of the RNA-seq screen. As measured by qPCR, PEG_2000_-PE/(GT)_6_-SWCNT suspensions induced either marginally or significantly lower upregulation of all 6 genes in SIM-A9 microglia compared to (GT)_6_-SWCNTs (Fig. 5D). In particular, *Cxcl2* expression change decreased significantly by 90 ± 2% when nanosensors were treated with PEG_2000_-PE. Upregulation of *Il1b* decreased by 47 ± 8%. Overall, SWCNT-induced expression changes of *Cxcl2, Il1b, Il6*, and *CCL4* were significantly lower than those induced by LPS.

Unlike covalent modification of the pristine carbon lattice surface, this passivation method preserved both the intrinsic SWCNT nIR fluorescence and the (GT)_6_-SWCNT molecular recognition for dopamine (fig. S7, A and B). Interestingly, the *in vitro* nanosensor response (ΔF/F_0_) upon addition of 200 μM dopamine increased upon (GT)_6_-SWCNT nanosensor passivation with PEG_2000_-PE at a 1:1 mass ratio, relative to the unpassivated (GT)_6_-SWCNT nanosensor, with ΔF/F_0_ = 2.01 and ΔF/F_0_ = 1.44, respectively. This effect was driven primarily by phospholipid-induced quenching of SWCNT baseline fluorescence. Furthermore, we tested whether PEG-passivated nanosensors would better withstand biofouling and nanosensor attenuation by blood plasma proteins. We found that the attenuation of (GT)_6_-SWCNT dopamine nanosensor response by plasma proteins was mitigated by PEG_2000_-PE passivation, where nanosensor incubation in 2% plasma caused nanosensor ΔF/F_0_ fluorescent response to decrease by 73% for unpassivated (GT)_6_-SWCNTs, compared to a 50% fluorescent response decrease for PEG_2000_-PE/(GT)_6_-SWCNTs (fig. S7C).

### Imaging striatal dopamine dynamics in acute mouse brain slice with passivated nanosensors

We imaged striatal dopamine release in acute mouse brain slices to evaluate the utility of PEG-phospholipid passivated SWCNT nanosensors as dopamine probes. PEG_2000_-PE/(GT)_6_-SWCNTs and (GT)_6_-SWCNTs were introduced into acute coronal brain slices, as previously described, by incubating fresh, 300 μm thick coronal brain slices in artificial cerebral spinal fluid (ACSF) containing 2 mg/L of dopamine nanosensor (Fig. 6A) (*11*). The nanosensor-labeled slices were then washed with ACSF and imaged in a continuously perfused ACSF bath. We electrically stimulated dopamine release from dopamine-containing axons within the dorsal lateral striatum and simultaneously imaged SWCNT nIR fluorescence response to changes in extracellular dopamine concentration. As expected, slices labeled with (GT)_6_-SWCNTs showed low nIR fluorescence signal prior to stimulation, followed by an increase in fluorescence response immediately after 0.3 mA electrical stimulation, and an eventual return to the low intensity - baseline ~5 s after stimulation (Fig. 6B). Brain slices labeled with PEG_2000_-PE/(GT)_6_-SWCNTs showed a similar nIR fluorescence response to 0.3 mA electrical stimulation (Fig. 6C), suggesting both the native dopamine probe and the PEG_2000_-PE-passivated probe enable imaging of dopamine release and re-uptake kinetics in brain tissue.

**Figure 6.**
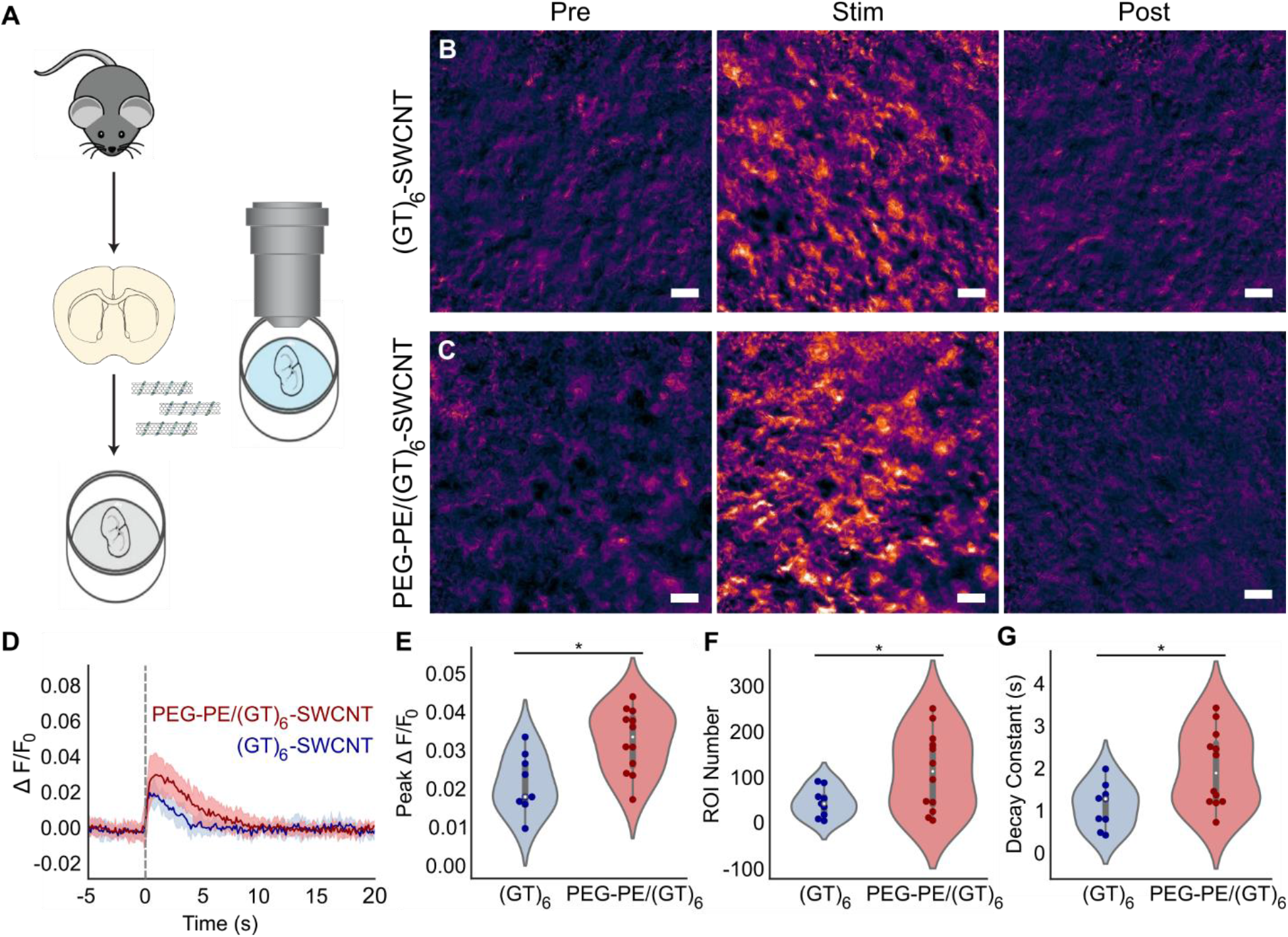
Imaging of dopamine release and reuptake dynamics in acute mouse striatal brain slices. **(A)** Schematic of acute mouse brain slice preparation and incubation with SWCNT nanosensors before dopamine release and reuptake imaging. **(B and C)** Representative images showing normalized nIR fluorescence signal (ΔF/F_0_) of **(B)** (GT)_6_-SWCNT and **(C)** PEG_2000_-PE/(GT)_6_-SWCNT in striatum of mouse brain before stimulation, at peak ΔF/F_0_ shortly after 0.3 mA single-pulse stimulation, and after SWCNT nanosensor signal returned to baseline. Scale bars are 10 μm. **(D)** Fluorescence response time trace of identified regions of interest (ROI) in brain slices labeled with (GT)_6_-SWCNT (blue) and PEG_2000_-PE/(GT)_6_-SWCNT during electrically evoked dopamine release. Dashed line indicates time of 0.3 mA single-pulse electrical stimulation. Solid lines represent mean traces and shaded regions represent one standard deviation around the mean for 4 mice, 1 brain slice per mouse, and 3 recordings per slice (N=12). **(E to G)** Violin plots showing the distribution of metrics from each mean nanosensor fluorescence trace for **(E)** peak ΔF/F_0_ signal, **(F)** number of identified regions of interest (ROIs), and **(G)** decay constant from fitting mean nanosensor ΔF/F_0_ time trace a first-order decay function. Dark points represent mean values calculated from each fluorescence video containing a single stimulation event. White dots represent the mean and the gray bar spans from the first to third quartiles. * p < 0.05.

We next characterized the spatial extent of nanosensor response to evoked dopamine release from striatal tissue. As described previously by Beyene *et al*. (*11*), we programmatically identified spatial regions of interest within the imaged brain tissue in which statistically significant increases in SWCNT fluorescence were recorded upon electrical stimulation (0.3 mA) of dopamine release. These regions of interest (ROI) represent spatial sub-regions where dopamine release and reuptake modulation occurs during electrical stimulation. Fluorescence time traces from ROIs were normalized to baseline fluorescence (ΔF/F_0_) and averaged across four brain slices per SWCNT treatment and three stimulation recordings per slice. Average ΔF/F_0_ of time traces from both (GT)_6_-SWCNT and PEG_2000_-PE/(GT)_6_-SWCNT labeled slices show that both nanosensors demonstrate a robust fluorescence response to dopamine released in living brain slices followed by a rapid return to baseline as dopamine is cleared from the extracellular space (Fig. 6D). For the same 0.3 mA stimulation intensity, PEG_2000_-PE/(GT)_6_-SWCNTs exhibited a peak ΔF/F_0_ of 0.032 ± 0.002 compared to 0.021 ± 0.003 for unmodified (GT)_6_-SWCNTs (Fig. 6E). This increased peak ΔF/F_0_ indicates improved dopamine responsivity by PEG_2000_-PE/(GT)_6_-SWCNTs compared to the unpassivated counterpart. PEG-phospholipid modified SWCNTs also improved ROI identification. In acute brain slices labeled with PEG_2000_-PE/(GT)_6_-SWCNTs, 158 ± 37 ROI were identified vs. 81 ± 15 ROI in (GT)_6_-SWCNT labeled slices (Fig. 5F). The higher ROI number may indicate improved extracellular access to dopaminergic terminals within the brain tissue. Conversely, PEG_2000_-PE/(GT)_6_-SWCNTs show significantly higher decay constants, indicating a slower return to baseline fluorescence (Fig. 6G). It is not known whether this effect is due to altered sensor kinetics arising from PEG-phospholipid modification or if it arises from the increased peak sensor ΔF/F_0_. As an additional control, stimulation at higher intensity (0.5 mA) revealed similar trends for the above metrics (fig. S8A). However, the increase in peak ΔF/F_0_ and ROI number from PEG_2000_-PE passivation was diminished (fig. S8, B and C). This may indicate saturation of the sensors from increased dopamine release at the higher electrical stimulation intensity. Nevertheless, the PEG_2000_-PE/(GT)_6_-SWCNT sensor displays higher sensitivity over (GT)_6_-SWCNT, particularly at lower analyte concentrations, suggesting dopamine nanosensors and other SWCNT-based neurotechnologies may benefit from this passivation approach.

## Conclusions

There have been numerous recent advances in the use of carbon nanomaterials for neuron stimulation, action potential recordings, brain morphological mapping, deep-brain imaging, and recording neuromodulatory kinetics. Herein, we first assess the impact of SWCNTs, a common carbon nanomaterial for above uses, on SIM-A9 microglial cells. Next, we present a passivation strategy that both preserves the dopamine response of the (GT)_6_-SWCNT nanosensor and mitigates the SWCNT-induced immune response. Microglial activation manifests in multiple cellular mechanisms, including a rapid change in cell morphology and upregulation of genes and pathways specific to the microglial immune response. We find that (GT)_6_-SWCNT nanosensors caused a large and persistent change in SIM-A9 morphology, transitioning from round, motile cells to multipolar, ramified cells with higher adhesion. This morphological effect was greater in magnitude than that induced by common immunogen LPS, and associated with extensive growth of actin cytoskeletal protrusions. The greater persistence and magnitude of morphology change induced by carbon nanotubes over LPS may be due to the relative persistence of SWCNTs within cellular environments, with degradation times on the order of days to weeks in tissue (*42, 43*). The full transcriptomic response induced by (GT)_6_-SWCNT nanosensors was also compared to other commonly used neuro-imaging and neuro-delivery probes, where the former uniquely showed a large change in the SIM-A9 transcriptomic profile.

Using high-throughput sequencing, we identified SIM-A9 genes that are highly upregulated in the presence of (GT)_6_-SWCNT nanosensors. We show that the predominant transcriptomic response to SWCNTs is an inflammatory response, whereby similarities in gene ontologies over-represented by differentially expressed genes from both (GT)_6_-SWCNT and LPS libraries suggest this effect is due to activation of toll-like receptors and the NF-κB signaling pathway. We note that the inflammatory response caused by 5 μg/mL (GT)_6_-SWCNTs is significantly lower in magnitude than that induced by a 1000-fold lower mass concentration of LPS. As such, the degree of neuroinflammation caused by SWCNTs is expected to be lower than that of a targeted pathogenic response.

To mitigate the immunological effects prompted by DNA-SWCNTs in microglia, we developed a noncovalent modification strategy to passivate the exposed SWCNT surface with PEGylated phospholipids. This methodology reduced protein adsorption by 28 ± 2%, and when applied to SIM-A9 microglia, resulted in a reduction of both inflammatory cytokine upregulation and a decrease in mean form factor change. This modification retains the SWCNT-based nanosensor nIR fluorescence response to dopamine and reduces attenuation of nanosensor signal by protein adsorption. Lastly, we apply the passivated PEG_2000_-PE/(GT)_6_-SWCNT nanosensor to image electrically-stimulated dopamine release and reuptake in acute mouse brain slices. Compared to unmodified (GT)_6_-SWCNTs, these nanosensors showed an increase in both fluorescence signal and responsivity in *ex vivo* mouse brain tissue increasing the max ΔF/F_0_ post stimulation by 52 ± 8%. Furthermore, PEG_2000_-PE/(GT)_6_-SWCNTs increased the number of identified ROI by 160 ± 50% potentially indicating improved dispersion of nanosensors or higher nanosensor sensitivity to dopamine release in tissue. Taken together, our data suggest that phospholipid PEG passivation of carbon nanotubes provides an avenue for improving both the biocompatibility and *in vivo* functionality of numerous SWCNT-based technologies already in proliferous use for neurobiological studies.

## Materials and Methods

### Preparation of neuro-sensors

Single-walled carbon nanotubes (SWCNTs) were dispersed in aqueous solution using (GT)_6_ single stranded DNA by combining 0.2 mg of small diameter HiPco™ SWCNTs (NanoIntegris) and 50 μM of ssDNA (Integrated DNA Technologies, Inc.) in 1 mL of 0.01 M phosphate-buffered saline (PBS). Solutions were probe-tip sonicated for 10 minutes using a 3 mm probe tip at 50% amplitude (5-6 W, Cole-Parmer Ultrasonic Processor). Following sonication, samples were centrifuged at 16,100 cfg for 30 minutes to pellet unsuspended SWCNT bundles, amorphous carbon, and metallic contaminants. Supernatant containing dispersed (GT)_6_-SWCNTs was collected. Excess DNA was removed via centrifugal filtration using an Amicon Ultra-0.5 mL centrifugal filter with a 100 kDa molecular weight cutoff (Millipore Sigma). Samples were placed in the filter and centrifuged at 8,000 cfg then washed with Milli-Q water. This process was repeated five times. Sample was recovered by reversing the spin filer and centrifuging into a collection tube at 1,000 cfg. Concentration of (GT)_6_-SWCNT suspensions was determined using sample absorbance at 632 nm and the corresponding extinction coefficient _ε632nm_ = 0.036 mL cm μg^-1^. (GT)_6_-SWCNTs were diluted to a 10x stock concentration of 100 μg/mL in 0.1 M PBS and stored at 4°C.

PEG-PE passivated (GT)_6_-SWCNTs were produced by mixing equal volumes of 200 μg/mL PEG-PE (Avanti Polar Lipids) and 200 μg/mL (GT)_6_-SWCNT in 0.1 M PBS. The mixture was bath sonicated for 15 min. Samples were used as prepared or stored at 4°C.

Carboxylic acid functionalized SWCNTs (Sigma Aldrich) were suspended in water by mixing approximately 1 mg solid COOH-SWCNT with 1 mL water and sonicating using the same settings detailed above. Centrifugation at 16,100 cfg for 30 minutes was again used to pellet and remove amorphous carbon, metallic catalysts, and unsuspended COOH-SWCNTs.

Prior to use in cell culture experiments, SWCNT suspensions were screened for endotoxin contamination using the Limulus amebocyte lysate (LAL) assay. Both (GT)_6_-SWCNT and COOH-SWCNT were confirmed to be below the limit of detection for endotoxin content.

Fura-2 AM, DiSBAC_2_(3), and Di-2-ANEPEQ (Thermo Fisher) were reconstituted in DMSO and diluted to a working concentration of 20 μM in PBS. rAAV1/Syn-GCaMP3 virus (UNC Vector Core, titer: 5×10^12^ virus molecules/mL) was diluted to a concentration of 2.5 x 10^11^ virus molecules/mL in PBS.

### Cell culture

Cyropreserved SIM-A9 cells were obtained from the UCB Cell Culture Facility and plated on a 75-cm^2^ culture flask in 10 mL of DMEM/F12 growth media supplemented with 10% fetal bovine serum, 5% horse serum, and 1x pen-strep-glutamine (Gibco, Life Technologies). All sera obtained were heat inactivated. Cells were stored in a humidified incubator at 37°C and 5% CO_2_. Cells were subcultured every 2-3 days after reaching approximately 90% confluence. Experiments were conducted using cells under passage number 15.

### Live-cell imaging

Cells were plated in a 96-well plate at a density of 50,000 cells per well in 100 μL of growth media. Cells were maintained at 37°C and 5% CO_2_ until approximately 70% confluent then washed with PBS. Media was replaced with sera free DMEM/F12 for two hours prior to start of experiments. Stock SWCNT or LPS was added to each well at 10x concentration, 0.1x total volume. Three biological replicates were run for each treatment. Phase contrast images were taken at 30-minute intervals using an IncuCyte^®^ Live-Cell Analysis System (Sartorius) in a humidified incubator at 37°C and 5% CO_2_. Images were analyzed using MATLAB (MathWorks) to identify and threshold cells for quantitation of cell area and perimeter.

### Confocal imaging of F-actin and DAPI stains

Cells were plated on poly-D lysine coated coverslips immersed in growth media in 6-well plates. Cells were treated with samples as previously described. Following treatment, cells were washed with PBS and fixed using 4% paraformaldehyde for 30 min at room temperature. Coverslips with fixed adherent cells were washed three times with PBS and submerged in 1 mL PBS. Two drops of ActinGreen 488 ReadyProbes Reagent (Thermo Fisher) were added and incubated for 1 h covered, at room temperature. DAPI counterstain was added to a final concentration of 1 μg/mL. The coverslip was rinsed three times and mounted in PBS on a glass microscope slide. Stained cells were imaged with a Zeiss LSM 710 laser scanning confocal microscope using DAPI and FAM fluorescence channels.

### RNA-seq library preparation and gene expression analysis

Cells were plated in a 24-well plate at a density of 0.1 x 10^6^ cells per well in 500 μL of growth media. Cells were maintained at 37°C and 5% CO_2_ until approximately 70% confluent then washed with PBS. Media was replaced with sera free DMEM/F12 for two hours prior to start of experiments. Stock neuro-sensor was added to wells at 10x concentration, 0.1x total volume. Three biological replicates were run for each group.

Two hours post exposure, total RNA was collected from adherent cells using the Quick RNA Miniprep Kit (Zymo Research) following manufacturer instructions. Cells were lysed directly on the plate and DNase treatment was used to remove genomic DNA. Total RNA concentration was measured using the Qubit™ RNA BR Assay Kit (Thermo Fisher). RNA quality was checked using the 2100 Bioanalyzer with RNA 6000 Nano Kit (Agilent). RIN scores were confirmed to be >7 prior to library preparation.

Libraries were prepared using Kapa Biosystems library preparation kit with mRNA selection with poly-A magnetic beads. Libraries were pooled and sequenced on an Illumina NovaSeq S4 flow cell with 150 paired end reads. Targeted data return was 25M read pairs per sample. Raw reads were pre-processed using *HTStream* (version 1.0.0) for filtering out adapter sequences, quality scores <30, and mouse ribosomal RNA (https://github.com/ibest/HTStream). Pre-processed reads were mapped to the Gencode M20 *Mus musculus* genome (GRCm38.p6) and quantified using *STAR* aligner (version 2.5.4b) (*44*). The *edgeR* package was used to determine differentially expressed genes (*33*). Adjusted p values were calculated using the Benjamini-Hockberg procedure using the *edgeR* default false discovery rate (FDR < 0.05). Gene ontology enrichment analysis was performed using the *topGO* R package (*34*). Enrichment of GO terms and p values were computed using Fisher’s exact test and the *weight01* algorithm with a p_adj_ < 0.01 cutoff for genes (*35*).

### Fluorescence tracking of protein adsorption

FAM fluorophore was conjugated to fibrinogen (FBG) using N-Hydroxysuccinimide (NHS) ester chemistry according to the protocol described in Pinals *et al* (*41*). SWCNT and FAM-FBG were mixed in a 1:1 volume ratio, 50 μL total in a 96 well PCR plate (Bio-Rad) and placed in a CFX96 Real-Time PCR System (Bio-Rad). Final concentrations were 5 μg/mL SWCNT and 40 μg/mL FAM-FBG. Scans were collected across all fluorescence channels (FAM, HEX, Texas Red, Cy5, Quasar 705) at 30 s intervals with temperature set to 22.5°C, lid heating off. A FAM-FBG fluorescence standard curve was used to convert fluorescence readings to unbound FAM-FBG concentrations.

### RT-qPCR

Total RNA from SIM-A9 cells with varying treatment conditions was reverse transcribed to cDNA libraries using the iScript cDNA Synthesis Kit (Bio-Rad) with a 1 μg RNA input. Next, 2 μL of cDNA was used with the PowerUp SYBR Green Master Mix (Thermo Fisher) and 500 nM of forward and reverse primers (table S2). The housekeeping genes, *Gadph* and *Pgk1* were used. Samples were cycled in a CFX96 Real-Time System (Bio-Rad) for 40 cycles (denature at 95°C for 15 s, anneal at 55° for 15 s, and extend at 72°C for 1 min). Data was analyzed using CFX Maestro software (Bio-Rad). Relative gene expression was calculated using the ΔΔCq method. P values were calculated using an unpaired t-test (N=3). RNA-seq experiments revealed that Pgk1 housekeeping gene did not undergo differential expression upon any treatment conditions. Melt curve analysis of RT-qPCR products was performed to ensure specific amplification.

### Mouse brain slice preparation and imaging

Acute brain slices were prepared from Male C57BL/6 Mice (JAX Strain 000664: https://www.jax.org/strain/000664) between the ages of 43-46 days. All mice were group-housed after weaning at postnatal day 21 (P21) and kept with nesting material on a 12:12 light cycle. All animal procedures were approved by the University of California Berkeley Animal Care and Use Committee. Preparation of acute brain slices followed previously established protocol (*11*). Mice were deeply anesthetized via intraperitoneal injection of ketamine/xylazine and perfused transcardially using ice-cold cutting buffer (119 mM NaCl, 26.2 mM NaHCO3, 2.5 mM KCl, 1 mM NaH2PO4, 3.5 mM MgCl2, 10 mM glucose, and 0 mM CaCl2). The perfused brain was extracted and the cerebellum removed. The brain was then mounted on to a vibratome (Leica VT1200 S) cutting stage using super glue and cut into 300 μm thick coronal slices containing the dorsal striatum. Slices were then transferred to 37°C oxygen-saturated ACSF (119 mM NaCl, 26.2 mM NaHCO3, 2.5 mM KCl, 1 mM NaH2PO4, 1.3 mM MgCl2, 10 mM glucose, and 2 mM CaCl2) for 30 minutes and then transferred to room temperature ACSF for 30 min. At this point, slices were ready for incubation and imaging and maintained at room temperature.

Prepared coronal slices were transferred into a small-volume incubation chamber (AutoMate Scientific) containing 5 mL oxygen-saturated ACSF. Two hundred microliters of 50 mg/L GT6-SWCNT or PEG_2000_-PE/(GT)_6_-SWCNT was added to the incubation chamber and the slice was allowed to incubate for 15 min. The slice was subsequently rinsed three times in baths of oxygen-saturated ACSF to wash off excess nanosensor. The labeled slice was transferred to an imaging chamber continually perfused with ACSF at 32°C. A bipolar stimulation electrode (125 μm Tungsten, 0.1 mΩ, WE3ST30.1A5 Micro Probes Inc.) was positioned in the dorsomedial striatum using a 4x objective (Olympus XLFluor 4x/340). The stimulation electrode was then brought into contact with the top surface of the brain slice 200 μm away from the imaging field of view using a 60x objective. All stimulation experiments were recorded at video frame rates of 9 frames per second for 600 frames and single, mono-phasic pulse (1 ms) electrical stimulations were applied after 200 frames of baseline were acquired. Each slice received pseudo-randomized stimulation at 0.3 mA and 0.5 mA, which were repeated three times each. Slices were allowed to recover for 5 min between each stimulation, with the excitation laser path shuttered. The excitation laser path was un-shuttered 1 min before beginning video acquisition.

Timeseries image stacks were processed using a suite of custom written MATLAB scripts (2019b MathWorks, https://github.com/jtdbod/Nanosensor-Brain-Imaging). A grid was superimposed over each frame to generate ~7 μm x 7 μm regions of interest (ROIs). For each ROI, the mean pixel intensity *F*(*t*) was calculated for each frame to generate an average intensity time trace. Δ*F*/*F*_0_(*t*) = (*F*(*t*) – *F*_0_)/*F*_0_ traces were generated with *F*_0_ calculated by averaging the mean ROI intensity of 10 frames prior to electrical stimulation followed by subtracting a linear offset to correct for drift. We estimated baseline noise (*σ*_0_) of Δ*F*/*F*_0_(*t*) by fitting a Gaussian to negative fluctuations from a moving averaged baseline. ROI’s were discarded from further analysis if no transient greater than 3σ_0_ was observed following stimulation. Remaining ROIs were then averaged to generate field of view (FOV) averaged ΔF/F_0_ traces for each recording.

From each mouse brain, two brain slices were selected at random and incubated with either (GT)_6_-SWCNTs or PEG_2000_-PE/(GT)_6_-SWCNTs. For each brain slice, three nIR fluorescence movies were collected at each stimulation intensity. A total of four mice brains were used, for a total of N=12 recordings per stimulation intensity, per nanosensor. Statistical analysis of metrics calculated from ΔF/F_0_ time traces was performed using an unpaired t-test to determine p values comparing (GT)_6_-SWCNT to PEG_2000_-PE/(GT)_6_-SWCNT incubated brain slices.

## Acknowledgements

RNA-seq libraries were prepared from extracted total RNA by the QB3 Genomics Functional Genomics Lab at UC Berkeley. Sequencing was performed by the Vincent J. Coates Genomics Sequencing Lab. Sequencing data are available in the Gene Expression Omnibus database under accession number GSE153419. M.P.L. acknowledges support of Burroughs Wellcome Fund Career Award at the Scientific Interface (CASI), Stanley Fahn PDF Junior Faculty Grant with Award #PF-JFA-1760, Beckman Foundation Young Investigator Award, DARPA Young Faculty Award, FFAR New Innovator Award, an IGI award, support from CITRIS and the Banatao Institute, and a USDA award. M.P.L. is a Chan Zuckerberg Biohub Investigator and an Innovative Genomics Institute Investigator. D.Y., S.J.Y., and R.L.P. acknowledge the support of the NSF Graduate Research Fellowship program. J.T.D.O. is supported by the Department of Defense office of the Congressionally Directed Medical Research Programs (CDMRP) Parkinson’s Research Program (PRP) Early Investigator Award.

## References

1. M. Kang, S. Jung, H. Zhang, T. Kang, H. Kang, Y. Yoo, J. P. Hong, J. P. Ahn, J. Kwak, D. Jeon, N. A. Kotov, B. Kim, Subcellular neural probes from single-crystal gold nanowires. ACS Nano. 8, 8182–8189 (2014).

2. A. C. Patil, N. V. Thakor, Implantable neurotechnologies: a review of micro- and nanoelectrodes for neural recording. Med. Biol. Eng. Comput. 54, 23–44 (2016).

3. S. Zhang, Y. Song, M. Wang, G. Xiao, F. Gao, Z. Li, G. Tao, P. Zhuang, F. Yue, P. Chan, X. Cai, Real-time simultaneous recording of electrophysiological activities and dopamine overflow in the deep brain nuclei of a nonhuman primate with Parkinson’s disease using nano-based microelectrode arrays. Microsystems Nanoeng. 4, 1–9 (2018).

4. E. A. Nance, G. F. Woodworth, K. A. Sailor, T. Y. Shih, Q. Xu, G. Swaminathan, D. Xiang, C. Eberhart, J. Hanes, A dense poly(ethylene glycol) coating improves penetration of large polymeric nanoparticles within brain tissue. Sci. Transl. Med. 4 (2012), doi:10.1126/scitranslmed.3003594.

5. H. Yang, Nanoparticle-mediated brain-specific drug delivery, imaging, and diagnosis. Pharm. Res. 27, 1759–1771 (2010).

6. S. Hrabetova, L. Cognet, D. A. Rusakov, U. V. Nägerl, Unveiling the extracellular space of the brain: From super-resolved microstructure to in vivo functiona. J. Neurosci. 38, 9355–9363 (2018).

7. E. W. Keefer, B. R. Botterman, M. I. Romero, A. F. Rossi, G. W. Gross, Carbon nanotube coating improves neuronal recordings. Nat. Nanotechnol. 3, 434–439 (2008).

8. A. C. Schmidt, X. Wang, Y. Zhu, L. A. Sombers, Carbon nanotube yarn electrodes for enhanced detection of neurotransmitter dynamics in live brain tissue. ACS Nano. 7, 7864–7873 (2013).

9. Z. Mohy-Ud-Din, S. H. Woo, J. H. Kim, J. H. Cho, Optoelectronic stimulation of the brain using carbon nanotubes. Ann. Biomed. Eng. 38, 3500–3508 (2010).

10. G. Hong, S. Diao, J. Chang, A. L. Antaris, C. Chen, B. Zhang, S. Zhao, D. N. Atochin, P. L. Huang, K. I. Andreasson, C. J. Kuo, H. Dai, Through-skull fluorescence imaging of the brain in a new near-infrared window. Nat. Photonics. 8, 723–730 (2014).

11. A. G. Beyene, K. Delevich, J. T. Del Bonis-O’Donnell, D. J. Piekarski, W. C. Lin, A. Wren Thomas, S. J. Yang, P. Kosillo, D. Yang, G. S. Prounis, L. Wilbrecht, M. P. Landry, Imaging striatal dopamine release using a nongenetically encoded near infrared fluorescent catecholamine nanosensor. Sci. Adv. 5, 1–12 (2019).

12. S. Jeong, D. Yang, A. G. Beyene, J. T. Del Bonis-O’Donnell, A. M. M. Gest, N. Navarro, X. Sun, M. P. Landry, High-throughput evolution of near-infrared serotonin nanosensors. Sci. Adv. 5, 1–13 (2019).

13. A. G. Beyene, K. Delevich, S. J. Yang, M. P. Landry, New Optical Probes Bring Dopamine to Light. Biochemistry. 57, 6379–6381 (2018).

14. A. G. Beyene, A. A. Alizadehmojarad, G. Dorlhiac, N. Goh, A. M. Streets, P. Král, L. Vuković, M. P. Landry, Ultralarge Modulation of Fluorescence by Neuromodulators in Carbon Nanotubes Functionalized with Self-Assembled Oligonucleotide Rings. Nano Lett. 18, 6995–7003 (2018).

15. C. Salvador-Morales, E. Flahaut, E. Sim, J. Sloan, M. L. H. Green, R. B. Sim, Complement activation and protein adsorption by carbon nanotubes. Mol. Immunol. 43, 193–201 (2006).

16. S. P. Mukherjee, O. Bondarenko, P. Kohonen, F. T. Andón, T. Brzicová, I. Gessner, S. Mathur, M. Bottini, P. Calligari, L. Stella, E. Kisin, A. Shvedova, R. Autio, H. Salminen-Mankonen, R. Lahesmaa, B. Fadeel, Macrophage sensing of single-walled carbon nanotubes via Toll-like receptors. Sci. Rep. 8, 1–17 (2018).

17. M. B. Graeber, W. J. Streit, Microglia: Biology and pathology. Acta Neuropathol. 119, 89–105 (2010).

18. D. J. Hines, R. M. Hines, S. J. Mulligan, B. A. Macvicar, Microglia processes block the spread of damage in the brain and require functional chloride channels. Glia. 57, 1610–1618 (2009).

19. G. Mukandala, R. Tynan, S. Lanigan, J. J. O’Connor, The effects of hypoxia and inflammation on synaptic signaling in the CNS. Brain Sci. 6 (2016), doi:10.3390/brainsci6010006.

20. M. T. Treadway, J. A. Cooper, A. H. Miller, Can’t or Won’t? Immunometabolic Constraints on Dopaminergic Drive. Trends Cogn. Sci. 23, 435–448 (2019).

21. J. C. Villegas, L. Álvarez-Montes, L. Rodríguez-Fernández, J. González, R. Valiente, M. L. Fanarraga, Multiwalled Carbon Nanotubes Hinder Microglia Function Interfering with Cell Migration and Phagocytosis. Adv. Healthc. Mater. 3, 424–432 (2014).

22. Y. Shigemoto-Mogami, K. Hoshikawa, A. Hirose, K. Sato, Phagocytosis-dependent and independent mechanisms underlie the microglial cell damage caused by carbon nanotube agglomerates. J. Toxicol. Sci. 41, 501–509 (2016).

23. K. Nagamoto-Combs, J. Kulas, C. K. Combs, A Novel Cell Line from Spontaneously Immortalized Murine Microglia. J NeurosciMethods, 759–785 (2014).

24. E. L. Gill, S. Raman, R. A. Yost, T. J. Garrett, V. Vedam-Mai, L -Carnitine Inhibits Lipopolysaccharide-Induced Nitric Oxide Production of SIM-A9 Microglia Cells. ACS Chem. Neurosci. 9, 901–905 (2018).

25. Q. Alhadidi, Z. A. Shah, Cofilin Mediates LPS-Induced Microglial Cell Activation and Associated Neurotoxicity Through Activation of NF-κB and JAK-STAT Pathway. Mol. Neurobiol. 55, 1676–1691 (2018).

26. T. Patriarchi, J. R. Cho, K. Merten, M. W. Howe, A. Marley, W. H. Xiong, R. W. Folk, G. J. Broussard, R. Liang, M. J. Jang, H. Zhong, D. Dombeck, M. von Zastrow, A. Nimmerjahn, V. Gradinaru, J. T. Williams, L. Tian, Ultrafast neuronal imaging of dopamine dynamics with designed genetically encoded sensors. Science (80-.). 360 (2018), doi:10.1126/science.aat4422.

27. H.-K. Jeong, K. Ji, K. Min, E.-H. Joe, Brain Inflammation and Microglia: Facts and Misconceptions. Exp. Neurobiol. 22, 59 (2013).

28. M. del M. Fernández-Arjona, J. M. Grondona, P. Fernández-Llebrez, M. D. López-Ávalos, Microglial Morphometric Parameters Correlate With the Expression Level of IL - 1β, and Allow Identifying Different Activated Morphotypes. Front. Cell. Neurosci. 13, 1–15 (2019).

29. L. P. Bernier, C. J. Bohlen, E. M. York, H. B. Choi, A. Kamyabi, L. Dissing-Olesen, J. K. Hefendehl, H. Y. Collins, B. Stevens, B. A. Barres, B. A. MacVicar, Nanoscale Surveillance of the Brain by Microglia via cAMP-Regulated Filopodia. Cell Rep. 27, 2895–2908.e4 (2019).

30. C.-M. Tîlmaciu, M. C. Morris, Carbon nanotube biosensors. Front. Chem. 3, 1–21 (2015).

31. X. Cui, B. Wan, Y. Yang, X. Ren, L. H. Guo, Length effects on the dynamic process of cellular uptake and exocytosis of single-walled carbon nanotubes in murine macrophage cells. Sci. Rep. 7, 1–13 (2017).

32. R. Parakalan, B. Jiang, B. Nimmi, M. Janani, M. Jayapal, J. Lu, S. S. W. Tay, E. A. Ling, S. T. Dheen, Transcriptome analysis of amoeboid and ramified microglia isolated from the corpus callosum of rat brain. BMC Neurosci. 13, 1 (2012).

33. M. D. Robinson, D. J. McCarthy, G. K. Smyth, edgeR: A Bioconductor package for differential expression analysis of digital gene expression data. Bioinformatics. 26, 139–140 (2009).

34. A. Alexa, J. Rahnenführer, topGO: Enrichment Analysis for Gene Ontology. R package (2019).

35. A. Alexa, J. Rahnenführer, T. Lengauer, Improved scoring of functional groups from gene expression data by decorrelating GO graph structure. Bioinformatics. 22, 1600–1607 (2006).

36. J. Zhao, H. Park, J. Han, J. P. Lu, Electronic Properties of Carbon Nanotubes with Covalent Sidewall Functionalization. J. Phys. Chem. B. 108, 4227–4230 (2004).

37. G. Bisker, J. Dong, H. D. Park, N. M. Iverson, J. Ahn, J. T. Nelson, M. P. Landry, S. Kruss, M. S. Strano, Protein-targeted corona phase molecular recognition. Nat. Commun. 7, 10241 (2016).

38. A. G. Godin, J. A. Varela, Z. Gao, N. Danné, J. P. Dupuis, B. Lounis, L. Groc, L. Cognet, Single-nanotube tracking reveals the nanoscale organization of the extracellular space in the live brain. Nat. Nanotechnol. 12, 238–243 (2017).

39. J. H. Choi, M. S. Strano, Solvatochromism in single-walled carbon nanotubes. Appl. Phys. Lett. 90, 88–91 (2007).

40. B. A. Larsen, P. Deria, J. M. Holt, I. N. Stanton, M. J. Heben, M. J. Therien, J. L. Blackburn, Effect of solvent polarity and electrophilicity on quantum yields and solvatochromic shifts of single-walled carbon nanotube photoluminescence. J. Am. Chem. Soc. 134, 12485–12491 (2012).

41. R. L. Pinals, D. Yang, A. Lui, W. Cao, M. P. Landry, Corona Exchange Dynamics on Carbon Nanotubes by Multiplexed Fluorescence Monitoring. J. Am. Chem. Soc. 142, 1254–1264 (2020).

42. M. Yang, M. Zhang, H. Nakajima, M. Yudasaka, S. Iijima, T. Okazaki, Time-dependent degradation of carbon nanotubes correlates with decreased reactive oxygen species generation in macrophages. Int. J. Nanomedicine. 14, 2797–2807 (2019).

43. V. E. Kagan, A. A. Kapralov, C. M. St. Croix, S. C. Watkins, E. R. Kisin, G. P. Kotchey, K. Balasubramanian, I. I. Vlasova, J. Yu, K. Kim, W. Seo, R. K. Mallampalli, A. Star, A. A. Shvedova, Lung macrophages Digest carbon nanotubes using a superoxide/peroxynitrite oxidative pathway. ACS Nano. 8, 5610–5621 (2014).

44. A. Dobin, C. A. Davis, F. Schlesinger, J. Drenkow, C. Zaleski, S. Jha, P. Batut, M. Chaisson, T. R. Gingeras, STAR: Ultrafast universal RNA-seq aligner. Bioinformatics. 29, 15–21 (2013).

