## Supplemental Information for "Transcriptomic and morphological response of SIM-A9 mouse microglia to carbon nanotube neuro-sensors"

#### S1. Supplementary Figures

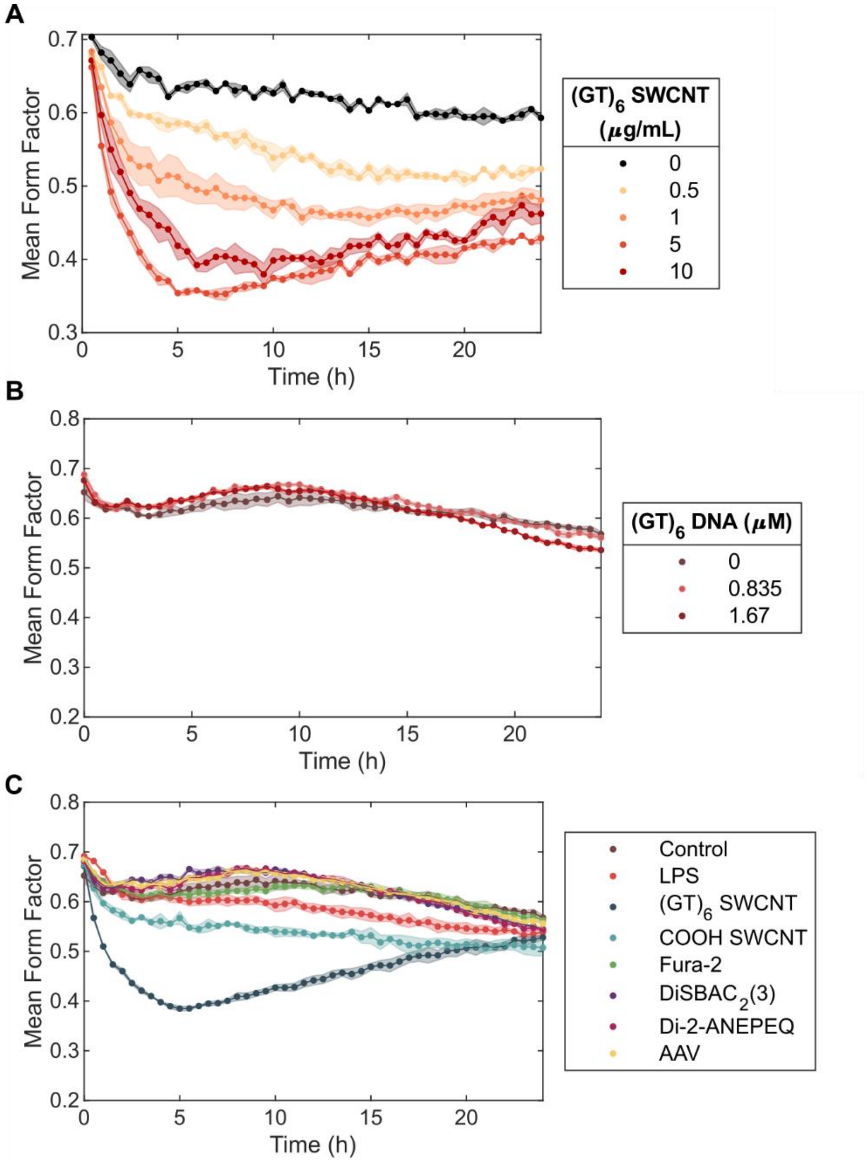


**Figure S1. SIM-A9 morphology change controls. (A)** Change in mean morphology upon addition of $(GT)_6$-SWCNT ranging in concentration from 0 to 10 μg/mL. SIM-A9 cells were at 37°C and 5% $CO_2$, in sera free media. **(B)** Change in mean cell form factor upon addition of free $(GT)_6$ ssDNA at concentrations equivalent to total ssDNA concentration in 5 and 10 μg/mL $(GT)_6$-SWCNT suspensions. **(C)** Cell morphology change upon addition of neurosensors at concentrations used for RNA-seq studies. SIM-A9 cells were at 37°C and 5% $CO_2$, in sera free media. Shaded regions represent standard error of the mean (N=3).

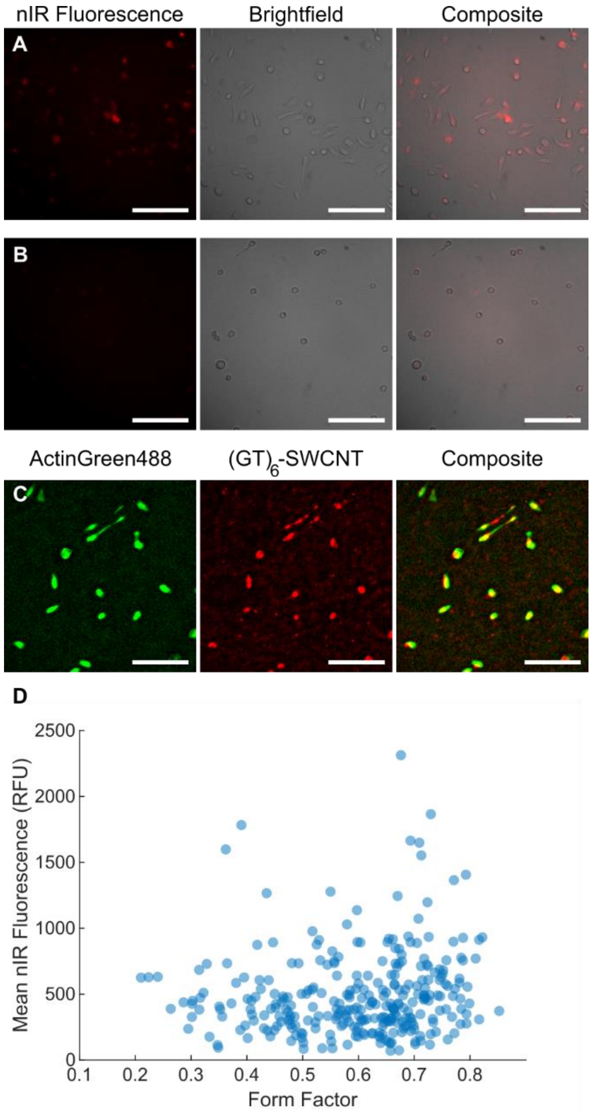


**Figure S2. Internalization of $(GT)_6$-SWCNTs in SIM-A9 microglia. (A-B)** SWCNT fluorescence, brightfield and composite images of fixed SIM-A9 microglia post incubation with 5 µg/mL $(GT)_6$-SWCNT for 1 hr at **(A)** 37°C and **(B)** 5°C. Scale bars are 50 µm. **(C)** Representative fluorescence images of ActinGreen488 stained SIM-A9 cells and internalized $(GT)_6$-SWCNTs following 2 h incubation at 37°C, 5% $CO_2$. Images are median filtered, background subtracted, and contrast adjusted. Scale bars are 50 µm. **(D)** Scatter plot of cell form factor versus the mean internalized nIR fluorescence signal. Form factor was calculated using the cell perimeter and area calculated from green fluorescence channel images of the stained actin cytoskeleton. A total of 302 cells were identified across 9 field of view captures of paired Alexa 488 and SWCNT fluorescence channels.

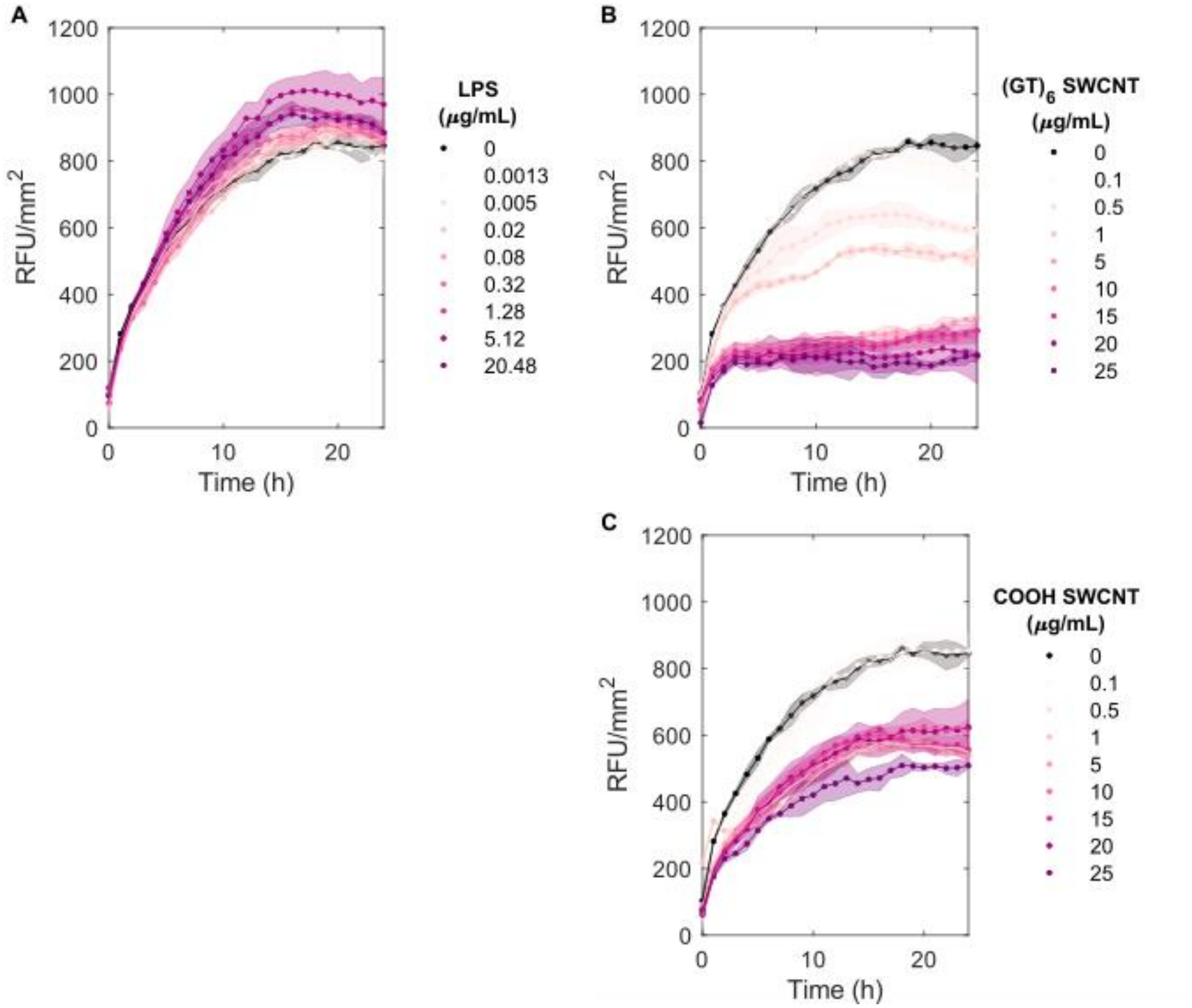


**Figure S3**. **SIM-A9 phagocytosis of fluorescent Zymosan bioparticles. (A-C)** Fluorescence live-cell imaging of pHrodo Red Zymosan Bioparticles (Sartorius) uptake by phagocytosis in SIM-A9 microglia following 3 h exposure to varying concentrations of **(A)** LPS, **(B)** $(GT)_6$-SWCNT, and **(C)** COOH-SWCNT. Shaded regions represent standard error of the mean (N=3).

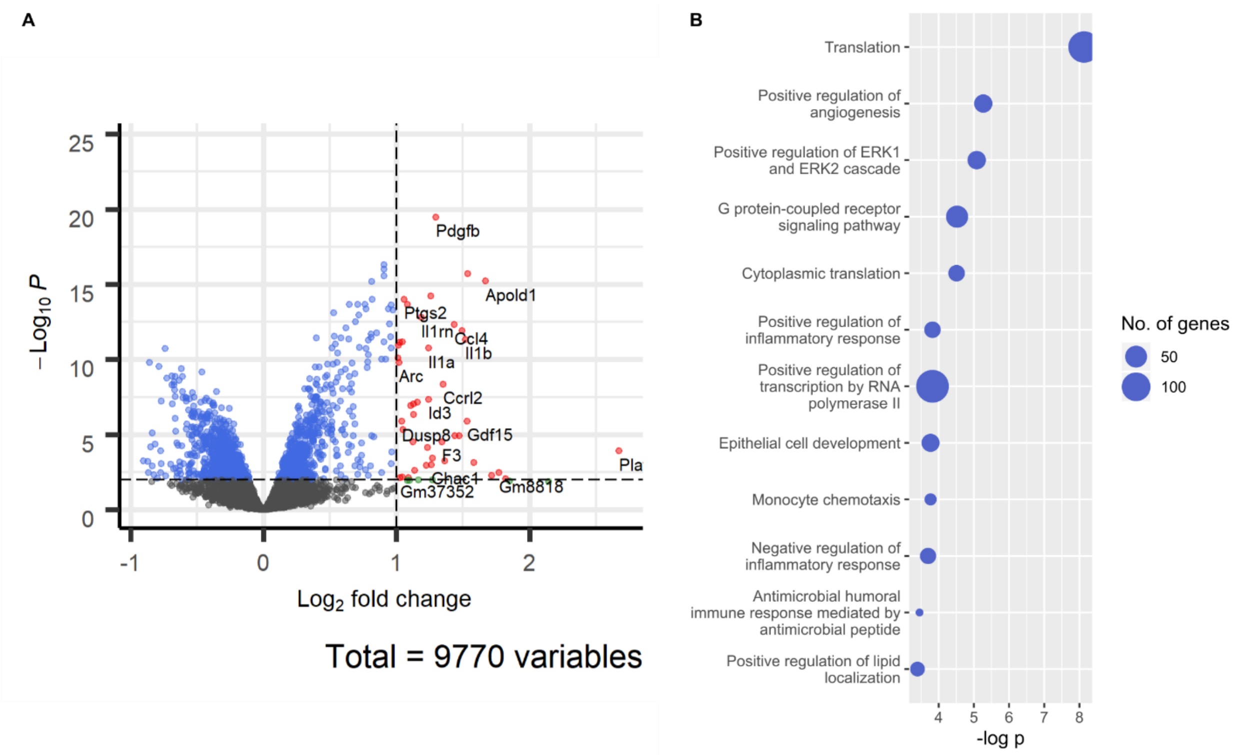


**Figure S4. COOH-SWCNT treated SIM-A9 differential gene expression. (A)** Volcano plot highlighting differentially expressed genes identified between COOH-SWCNT and untreated cell control libraries. **(B)** Top 12 gene ontologies overrepresented by DE genes in COOH-SWCNT incubated microglia.

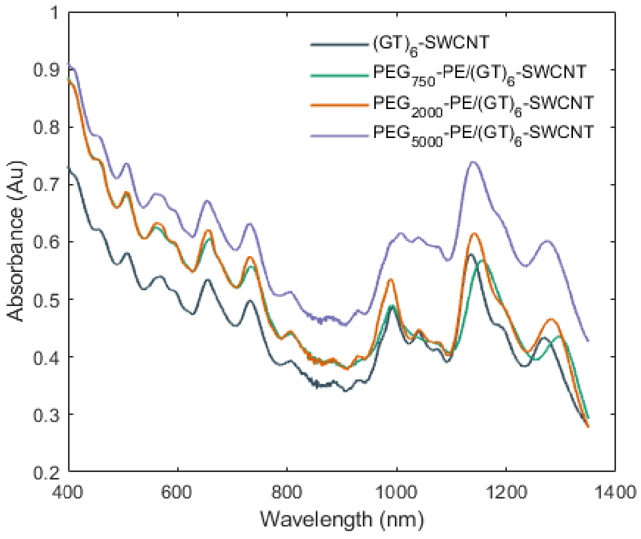


**Figure S5. Absorbance spectra of PEG-PE passivated $(GT)_6$-SWCNTs.** UV-vis-nIR absorbance of 20 μg/mL $(GT)_6$-SWCNT, $PEG_{750}$-PE/$(GT)_6$-SWCNT, $PEG_{2000}$-PE/$(GT)_6$-SWCNT, and $PEG_{5000}$-PE/$(GT)_6$-SWCNT. Spectra collected on a UV-3600i Plus UV-Vis-NIR Spectrophotometer using dual beam measurement with stair correction.

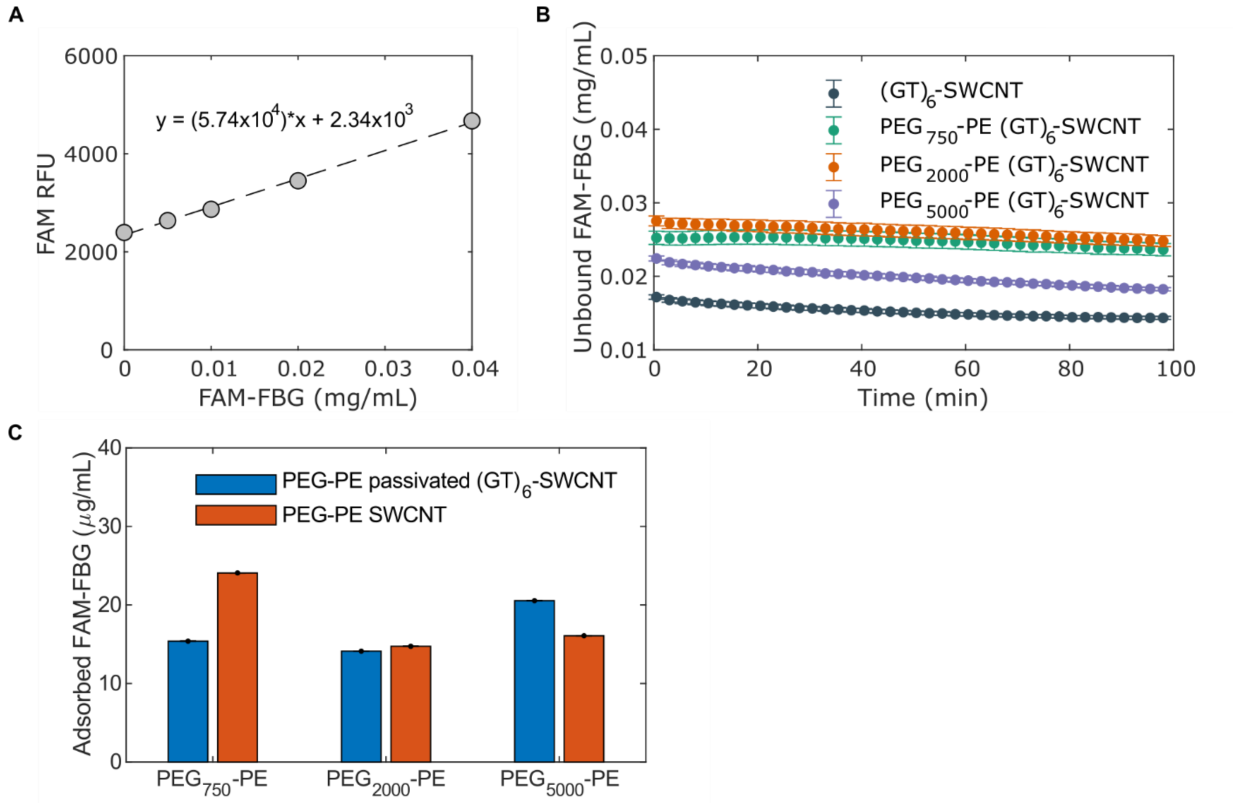


**Figure S6. Quantification of real-time FAM-fibrinogen adsorption to SWCNTs. (A)** FAM-FBG fluorescence calibration curve. **(B)** Concentration time series of unbound FAM-FBG. Protein added at total concentration of 40 μg/mL FAM-FBG to 5 μg/mL SWCNT. **(C)** Adsorption of 40 μg/mL FAM-FBG determined by quenching of conjugated FAM fluorophore to 5 μg/mL $(GT)_6$-SWCNTs passivated with PEG-PE of varying PEG molecular weights vs. SWCNTs suspended with PEG-PE.

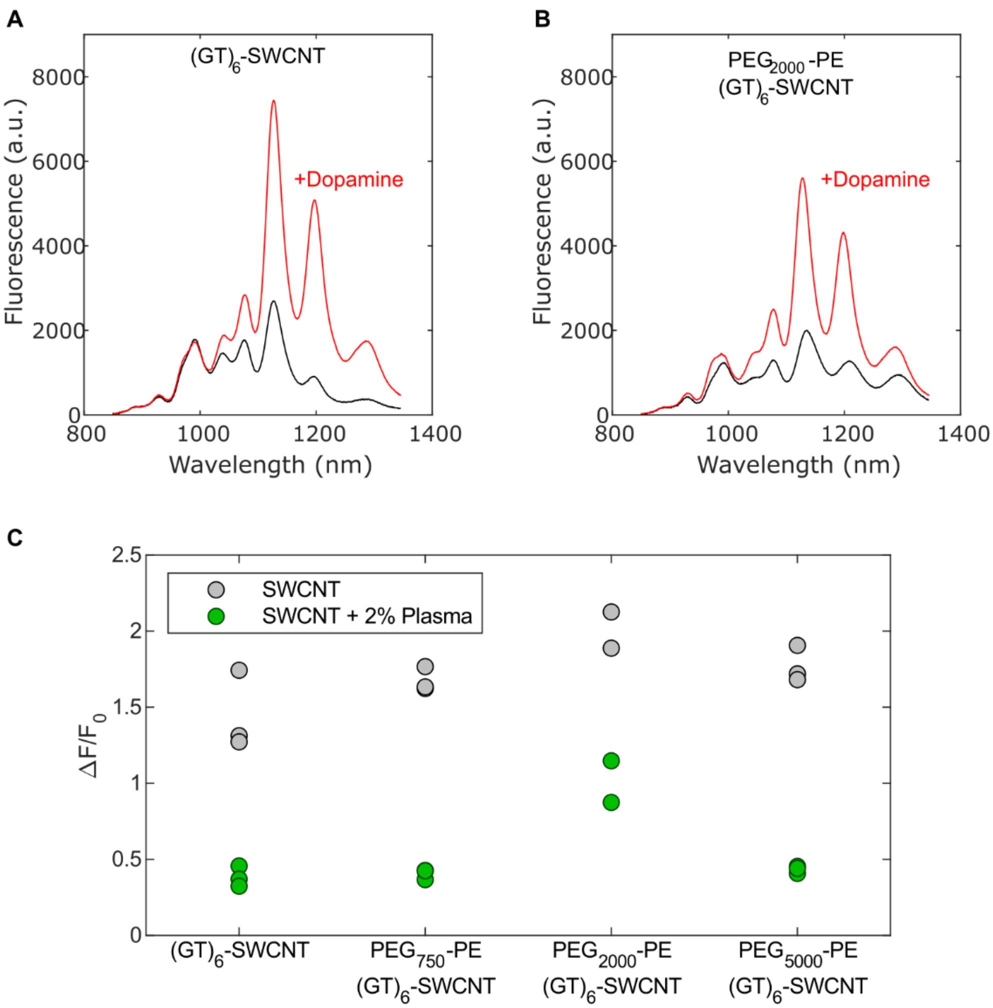


**Figure S7. Dopamine response of PEG-PE passivated SWCNTs. (A-B)** Near infrared fluorescence spectra of **(A)** $(GT)_6$-SWCNT and **(B)** $PEG_{2000}$-PE passivated $(GT)_6$-SWCNT before and after addition of 200 µM dopamine. **(C)** Fluorescence response to 200 µM dopamine of 5 µg/mL $(GT)_6$-SWCNT and PEG-PE/$(GT)_6$-SWCNT with PEG molecular weights of 750, 2000, and 5000 Da (grey). Dopamine response in biological milieu was simulated by pre-incubating SWCNT suspensions with 2% human blood plasma for 15 min (green).

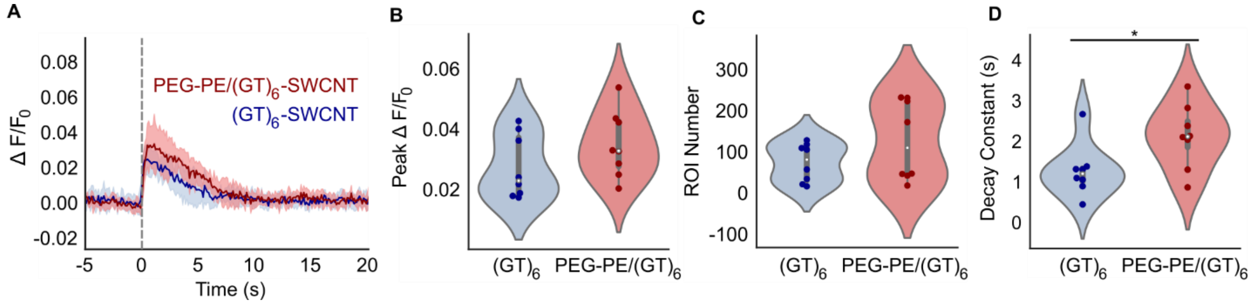


**Figure S8**. **Striatal dopamine release in acute mouse brain slice with 0.5 mA electrical stimulation. (A)** Near infrared fluorescence time traces of regions of interest (ROI) identified in acute mouse brain slice labeled with $(GT)_6$-SWCNTs (blue) and $PEG_{2000}$-PE/$(GT)_6$-SWCNTs (red). Dashed line indicates 0.5 mA single-pulse electrical stimulation. Solid lines represent mean traces and shaded regions represent one standard deviation around the mean for 4 mice, 1 brain slice per mouse, and 3 recordings per slice (N=12). **(B-D)** Violin plots showing the distribution of metrics from each mean nanosensor fluorescence trace for **(B)** peak $\Delta F/F_0$, **(C)** number of identified regions of interest (ROIs), and **(D)** decay constant from fitting mean nanosensor $\Delta F/F_0$ time trace a first-order decay function. Dark points represent mean values calculated from each fluorescence video containing a single stimulation event. White dots represent the mean and the gray bar spans from the first to third quartiles. * $p < 0.05$.

**Table S1. Summary of differentially expressed genes identified in LPS, $(GT)_6$-SWCNT, and COOH-SWCNT stimulated microglia vs. untreated control cells.**

| | | **LPS vs. Control** | | | **$(GT)_6$-SWCNT vs. Control** | | | **COOH-SWCNT vs. Control** | | |
|---|---|---|---|---|---|---|---|---|---|---|
| | | Up | Down | Total | Up | Down | Total | Up | Down | Total |
| DE Genes | $p_{adj} < 0.05$ | 1558 | 1706 | 3264 | 1204 | 1242 | 2446 | 1107 | 1346 | 2453 |
| | $log_2$ Fold Change > 1 | 239 | 93 | 332 | 105 | 14 | 119 | 49 | 0 | 49 |
| Gene Type | Protein Coding | 1481 | 1663 | 3144 | 1149 | 1218 | 2367 | 1066 | 1278 | 2344 |
| | Long noncoding RNA | 50 | 34 | 84 | 37 | 16 | 53 | 24 | 37 | 61 |
| | Pseudogene | 15 | 4 | 19 | 14 | 5 | 19 | 9 | 26 | 35 |

**Table S2. Primers for RT-qPCR**

| **Gene** | **Forward Strand (5' -> 3')** | **Reverse Strand (5' -> 3')** |
|---|---|---|
| GADPH | ACCACAGTCCATGCCATCAC | CTGGTTCCTGAAGCGACAAC |
| Pgk1 | CTGACTTTGGACAAGCTGGAC | GCAGCCTTGATCCTTTGGTTG |
| Cxcl2 | CTCTCAAGGGCGGTCAAAAAGTT | TCAGACAGCGAGGCACATC |
| IL-1b | GCACTACAGGCTCCGAGATGA | TTGTCGTTGCTTGGTTCTCCTTG |
| IL6 | CCTCTGGTCTTCTGGAGTACC | ACTCCTTCTGTGACTCCAGC |
| PDGFb | GTTGCAACGAGAAAGCCGGAG | GTCTGTCTATCTACCCACTCGC |
| CCL4 | CCCAGCTCTGTGCAAACCTA | CCATTGGTGCTGAGAACCCT |
| IL-1b | TGCCACCTTTTGACAGTGATG | AAGGTCCACGGGAAAGACAC |

**Movie S1. Live-cell imaging time lapse video of untreated SIM-A9 microglia**

**Movie S2. Time lapse video of SIM-A9 microglia treated with 5 µg/mL $(GT)_6$-SWCNT**

#### S2. SI Materials and Methods

##### SWCNT internalization into SIM-A9

Cells were plated in 35mm dishes at a density of 0.15 x $10^6$ cells per well in growth media and incubated overnight. Cells were serum starved for two hours prior to imaging. Concentrated $(GT)_6$-SWCNTs in PBS was added to a final concentration of 5 µg/mL. Following incubation under specified conditions, media was removed and cells were washed with PBS and subsequently fixed with 4% PFA incubation for 20 min at room temperature. Fixed cells were washed with PBS twice then stained with ActinGreen 488 (Invitrogen) for 30 min at room temperature. Cells were washed with PBS before imaging on a Zeiss upright microscope (Axio Observer.D1) with a 10x objective. A 721 nm laser was used for excitation and signal was collected with a Ninox 640 InGaAs camera (Raptor Photonics). Brightfield images were collected using the same camera with LED illumination. Green fluorescence images were collected using LED illumination and a FITC filter set (Chroma).

##### Phagocytosis assay

SIM-A9 microglia were plated on a 96 well plate at a density of 10,000 cells per well. After overnight incubation in a humidified incubator at 37°C and 5% $CO_2$ media was replaced with 90 µL sera free DMEM/F12 and incubated for an additional 4 hours. A 10 µL aliquot of sample at 10x concentration (LPS, $(GT)_6$-SWCNT, COOH-SWCNT or PBS control) was added to each corresponding well. Following 3 h incubation at 37°C and 5% $CO_2$, a 5 µL aliquot of 1 mg/mL pHrodo red (Sartorius) was added to each well. The 96 well plate was imaged in an IncuCyte® Live-Cell Analysis System (Sartorius) using the phase contrast and red fluorescence channels at 1 h intervals. Mean fluorescence per cell area was computed using the IncuCyte Base Analysis software.

##### Near infrared fluorescence measurements

SWCNTs were diluted to a concentration of 5 µg/mL in PBS. A 40 µL aliquot was added to a 384 well plate with optical bottom (Corning). The plate was placed on a motorized stage of an inverted Zeiss microscope (Axio Observer.D1) coupled to an InGaAs array detector (Princeton Instruments). SWCNT nIR fluorescence spectra were measured using a 10x objective with 721 nm laser excitation. Fluorescence response to dopamine was measured by collecting spectra before and after addition of 10 µL, 100 µM dopamine hydrochloride.

##### Limulus amebocyte lysate (LAL) assay for endotoxin contamination

The Pierce Chromogenic Endotoxin Quant Kit (Thermo Scientific) was used to determine endotoxin contamination in SWCNT samples. These samples were diluted to 0.3 μg/mL, the maximum concentration before SWCNTs begin to induce false activation of the enzymatic reaction utilized.[1] LPS was diluted to 0.3 ng/mL to provide a direct comparison to concentrations used in RNA-seq, and to 0.05 ng/mL. Samples were vortexed rigorously in order to detach endotoxins adsorbed the tube sides. The LAL assay was run according to manufacturer instructions with each sample run in experimental triplicate. SWCNT samples were confirmed to contain endotoxin concentrations below the LAL assay limit of detection, 0.1 EU/mL where 1 endotoxin unit (EU) corresponds to approximately 0.1 – 0.2 ng of endotoxin equivalent.

### SI References

(1) Yang, M.; Nie, X.; Meng, J.; Liu, J.; Sun, Z.; Xu, H. Carbon Nanotubes Activate Limulus Amebocyte Lysate Coagulation by Interface Adsorption. *ACS Appl. Mater. Interfaces* **2017**, *9* (10), 8450–8454. https://doi.org/10.1021/acsami.7b00543.
